# SLFN11 Enhances Cisplatin Sensitivity in Pediatric Cancer via Activation of Stress-Response and Suppression of Survival Pathways

**DOI:** 10.1101/2025.06.17.659735

**Authors:** Ayesha Jabeen, Dina Awartani, Shimaa Sherif, Eiman I. Ahmed, Rania Alanany, Ayman Saleh, Wouter R.L. Hendrickx, Christophe M. Raynaud

**Author notes:** These author contributed equally to this work (Ayesha Jabeen, Dina Awartani). Corresponding authors: Christophe Michel Raynaud, Wouter R.L. Hendrickx.

## Abstract

Pediatric cancers pose significant treatment challenges due to their biological heterogeneity and variable responses to chemotherapy. SLFN11, a DNA/RNA helicase-like protein known to sensitize adult tumors to DNA-damaging agents, remains underexplored in pediatric malignancies. Here, we investigate the role of SLFN11 across pediatric Wilms tumor, osteosarcoma, and medulloblastoma using integrated bioinformatics, epigenetic profiling, and functional assays. In silico analysis of TARGET and ICGC datasets revealed distinct correlations between SLFN11 expression and patient survival, with positive, negative, or neutral predictive value depending on tumor type. Baseline expression and promoter methylation analysis in pediatric cancer cell lines demonstrated epigenetic regulation of SLFN11, similar to adult cancers. Using CRISPR-dCas9-mediated activation, we successfully upregulated SLFN11, which significantly enhanced sensitivity to cisplatin and the PARP inhibitor talazoparib across all tested cell lines. Transcriptomic profiling under cisplatin treatment indicated that SLFN11 modulates DNA damage response and MAPK signaling pathways, potentially contributing to chemotherapy sensitivity. These findings establish SLFN11 as a context-dependent predictive biomarker and a potential therapeutic target to overcome chemoresistance in pediatric solid cancers.

## Introduction

SLFN11, initially identified as a putative DNA/RNA helicase localized in the nucleus, gained prominence in 2012 when its expression was linked to sensitivity to DNA-damaging agents (DDAs), including platinum-based drugs like cisplatin and PARP inhibitors such as talazoparib [1]. Since this seminal work SLFN11 was extensively studied. During replication stress, SLFN11 binds to Replication Protein A (RPA) at stalled replication forks and interacts with MCM3 and DHX9. These interactions facilitate chromatin remodeling around replication initiation sites, activating the transcription of immediate early genes and inducing cell cycle arrest [2,3], leading to the opening of the chromatin around the replication initiation sites, which activates the transcription of immediate early genes that can induce cell cycle arrest. Thereby blocking any further replications from occurring [4]. This is done independently of, and in parallel with, the ATR-CHEK 1 S-phase checkpoint in the DDR pathway [2]. Given the frequent defects in the DDR pathway observed in cancers [5], SLFN11 may serve as a critical fail-safe mechanism for inducing cell cycle arrest in response to DNA damage.

SLFN11 expression is tightly regulated at the epigenetic and transcriptional levels. In cancer cells, SLFN11 is frequently silenced through promoter methylation, resulting in resistance to DNA-damaging agents and PARP inhibitors [2,3,6,7]. In a study of 66 human SCLC cell lines, higher promoter methylation correlated with reduced SLFN11 expression and treatment resistance [7]. Three main mechanisms suppress SLFN11 gene expression: promoter methylation, histone methylation by the Polycomb Repressive Complex (PRC), and histone deacetylation [2,3,8,9].

SLFN11 is also regulated by interferon-gamma (IFN-γ), linking it to the innate immune response. IFN-γ, crucial for immune defense, induces SLFN11, which can predict sensitivity to treatments like Olaparib and Temozolomide in small cell lung cancer (SCLC) [10–13]. Additionally, SLFN11 induction during viral infections, such as Zika virus or HIV, correlates with viral inhibition, suggesting a shared pathway between innate immune activation and DDA response [14].

While most studies on SLFN11 have focused on adult cancers, our investigation explores its role as a predictive biomarker in pediatric cancers, which often exhibit distinct genetic and molecular characteristics. Conflicting results were reported in the literature regarding SLFN11 and response to DNA Damage agents. For instance, in Ewing sarcoma, SLFN11 is upregulated by the EWS-FLI1 fusion and correlates with better response to PARP inhibitors and improved survival [15]. In medulloblastoma, especially WNT and SHH subtypes, high SLFN11 enhances cisplatin sensitivity and can be upregulated with HDAC inhibitors [16]. A pan-cancer analysis showed SLFN11 expression in 69% of pediatric solid tumors, notably in Ewing sarcoma (90%) and desmoplastic small round-cell tumors (100%). While SLFN11 predicts DDA response in preclinical models, clinical outcomes suggest additional factors may influence its predictive value.[17].

We aim to understand the activity of SLFN11 in pediatric tumors, and whether it is similar or not to its role in adult tumors. The understanding SLFN11’s function in these cancers could reveal unique mechanisms of treatment sensitivity or resistance.

Understanding how SLFN11 expression and function differ in pediatric tumors could not only influence prediction of treatment outcomes but also pave the way for more personalized therapeutic strategies. Such advancements hold the potential to improve survival rates and minimize treatment-related side effects for pediatric cancer patients.

## Material and method

### Cell lines and culture

All cell lines were purchased from ATCC (**Table 1**). All cell lines were adapted and cultured in advanced RPMI (Gibco, #12633012) complemented with 10% FBS (Gibco, #10500-064), Glutamax (Gibco, #35050061) and antibiotic-antimycotic (Gibco, #15240096). Cells were cultured at 37°C, 5% CO2 and 95% humidity. Cells were detached using TrypLE express enzyme (Gibco, #12605036). Human foreskin fibroblasts were used across the manuscript as positive control and reference for expression of SLFN11.

**Table 1:**
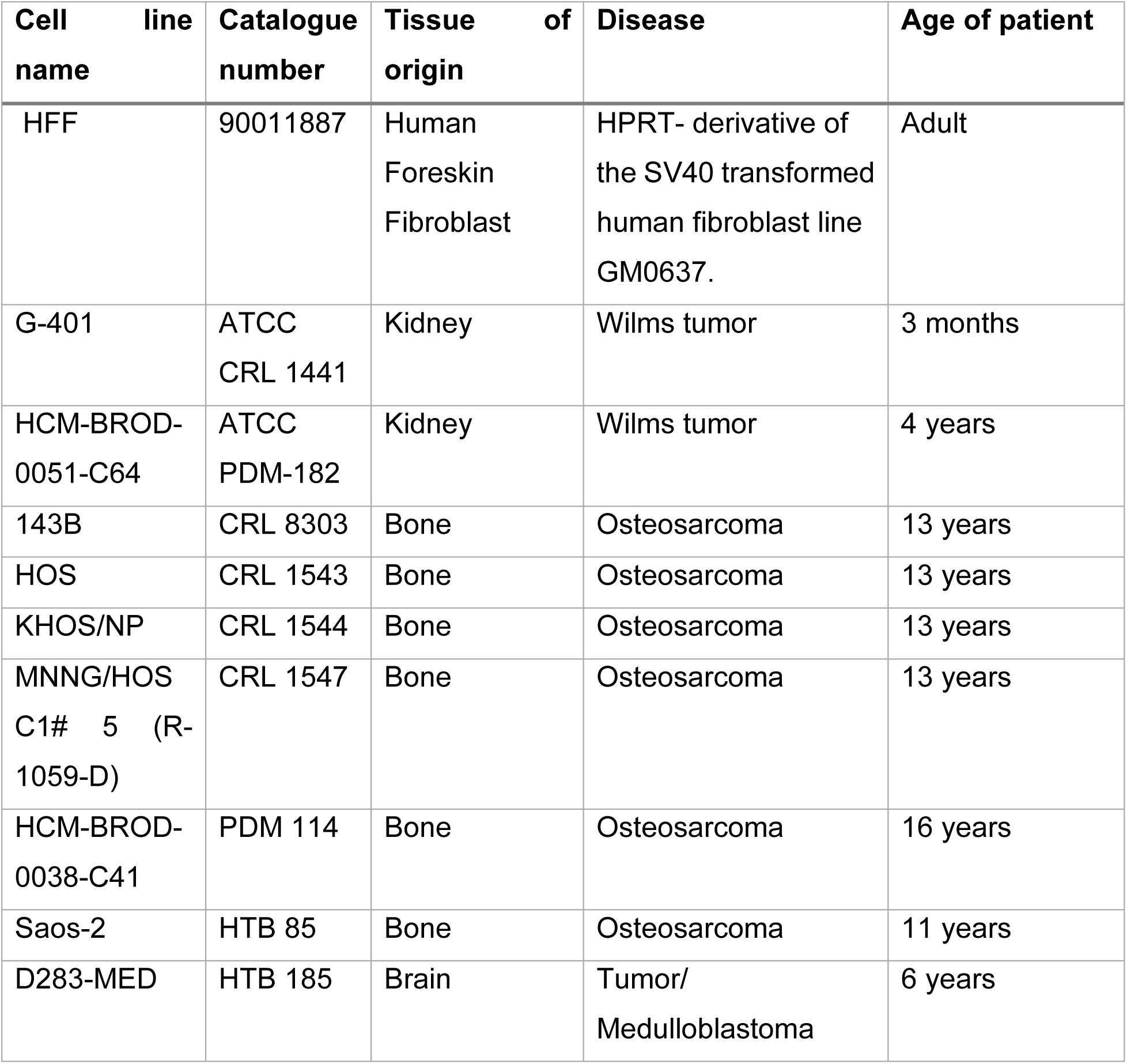
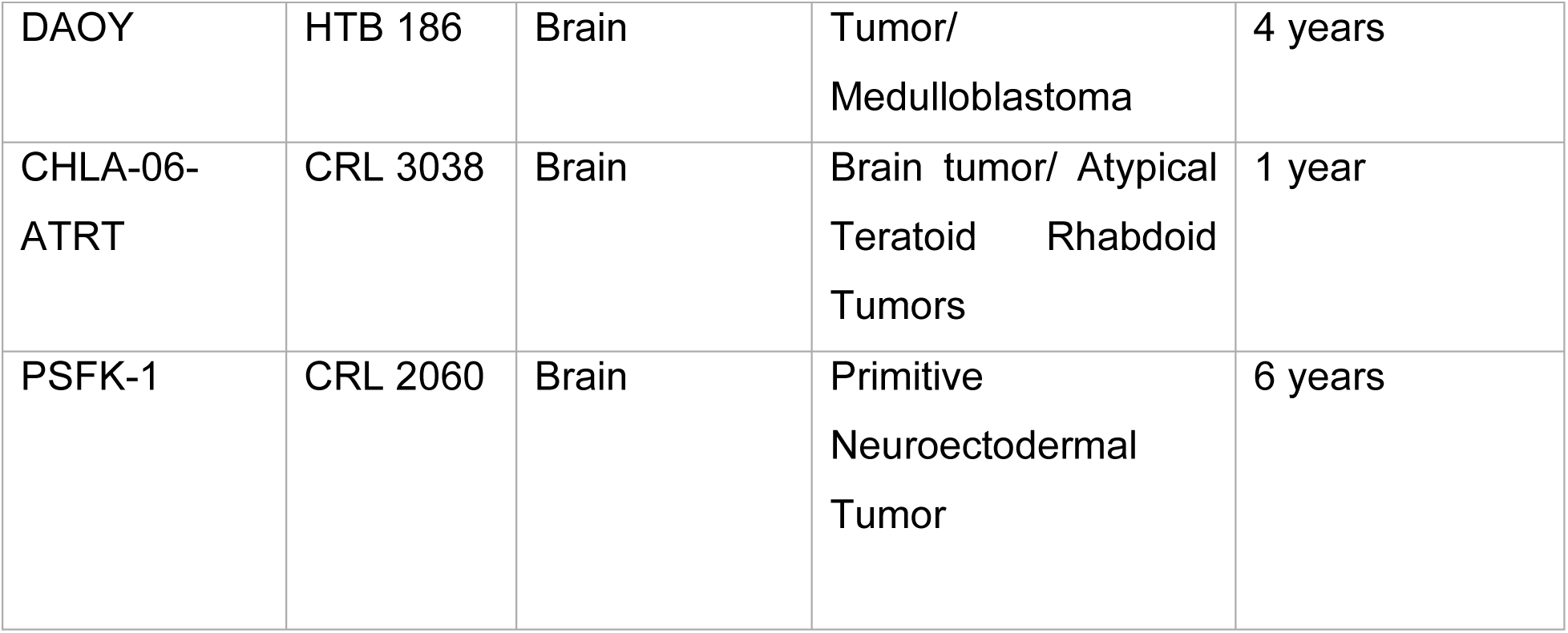
cell lines.

### CRISPR cell engineering - gRNA design

gRNAs were previously designed [18] using IDT custom gRNA design tool along the core region of the promoter,

The sequence of the gRNA used is provided in **table 2**.

**Table 2:**
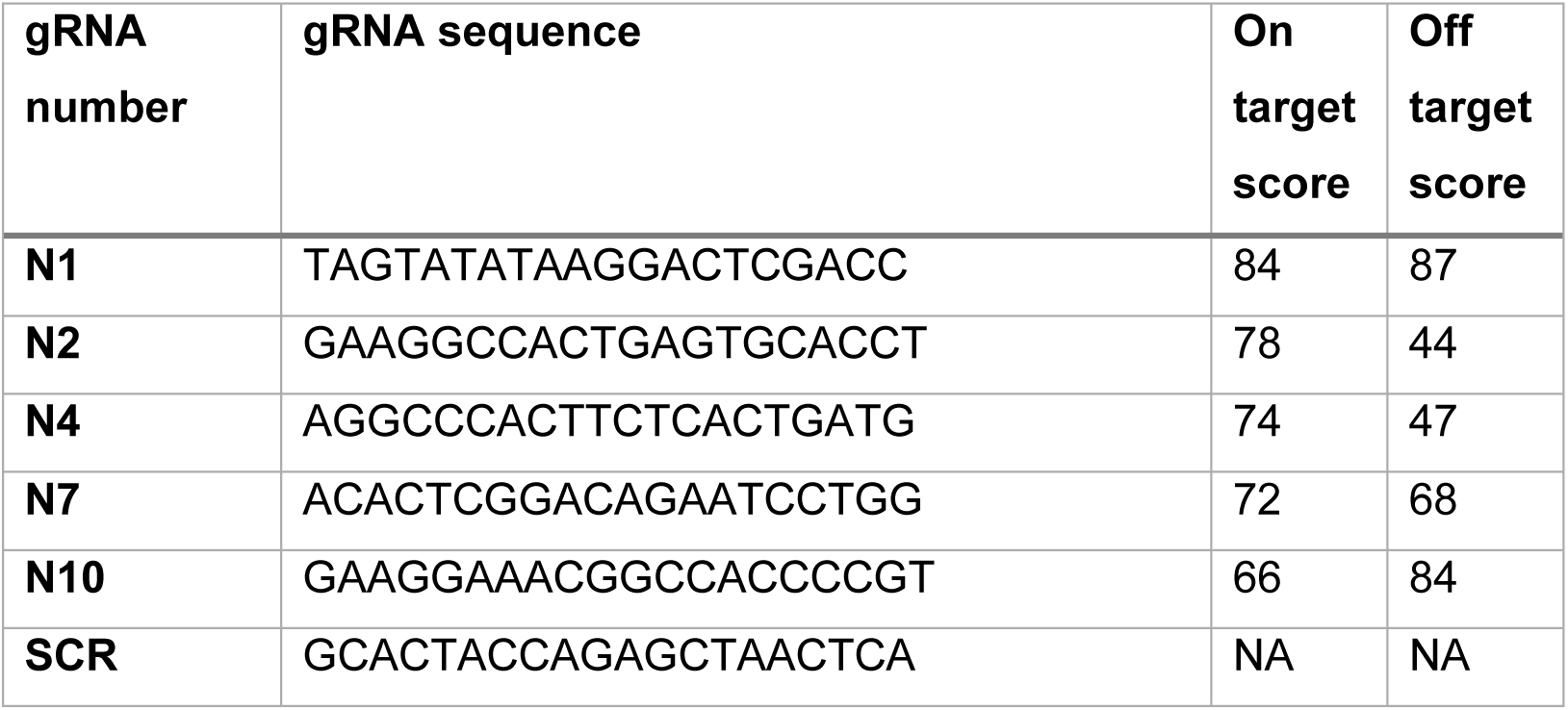
gRNA sequence.

The sequence of the scramble (SCR) gRNA used is GCACTACCAGAGCTAACTCA. Forward and reverse primers were ordered accordingly to be inserted in the appropriate plasmids as described below.

### DNA/RNA extraction

DNA isolation was performed using Maxwell® RSC Cultured Cells DNA Kit (Promega, # AS1620) according to manufacturer recommendations using Maxwell® RSC Instrument (Promega, #AS4500). DNA was recovered with 100ul of elution buffer and measured with QuantiFluor® ONE dsDNA System (Promega, # E4871). RNA was extracted Maxwell® RSC simplyRNA Cells Kit (Promega, #AS1390) according to manufacturer recommendations using Maxwell® RSC Instrument (Promega, #AS4500). RNA was recovered with 50ul of RNase free water and measured with QuantiFluor® RNA System (Promega, #E3310). DNA and RNA were stored at −80°C until use.

### Q-PCR

1ug of RNA was used for reverse transcription using TaqMan™ Reverse Transcription Reagents (Invitrogen, #N8080234) and random hexamers. cDNA was diluted 20 times with DNA/RNA free water. TaqMan™ Gene Expression Master Mix (Applied bioscience, #4369016) and Hs03003631_g1 (for Eukaryotic 18S rRNA) and Hs00536981_m1 (for SLFN11) TaqMan probes (Thermo scientific, #4331182) were used according to manufacturer recommendation. Quantitative Real time PCR (Q-RT-PCR) was run in 96 well plates on QuantStudio 12K flex system (Thermofisher Scientific). Q-RT-PCR was done in triplicate for each sample and data were analyzed by gene expression comparison using ΔΔCT on (QuantStudio 12K Flex Realtime PCR system V1.2.2) using S18 as the housekeeping gene.

### CWB

Capillary western blot (CWB) was done using a WES system (protein simple) with 12-230 kDa Separation Module, 8 x 25 capillary cartridges (Protein simple, #SW-W004), EZ Standard Pack 2 (Protein simple, #PS-ST02EZ-8) and Anti-mouse detection module (Protein simple DM-002). Mouse anti human SLFN11 (Santa Cruz, #SC-515071) and anti β-actin (Licor, #926-42212) both diluted at 1 in 100 were used as primary antibody.

Analysis was done using compass for Simple western (ProteinSimple, V5.0.0) and area of histogram peaks were used for quantification. All western blot analysis were normalized for β-actin expression.

### Promoter Methylation Analysis by MSP

Promoter methylation was analyzed using Methyl Specific Polymerase Chain Reaction (MSP). Genomic DNA was extracted as previously described, bisulfite conversion was performed using EZ DNA methylation kit (Zymo research, # D5001). PCR was performed using the primers: Forward Methylated specific primer (GTAGCGGGGTAGAAAAGTAGAAC) and Reverse Methylated specific primer (TAAAATTTAACGACGACCGATACG) for methylated specific PCR with a PCR product of 108bp. Forward Unmethylated specific primer (GTAGTGGGGTAGAAAAGTAGAAT) and Reverse unmethylated specific primer (TAAAATTTAACAACAACCAATACA) for unmethylated specific PCR with a PCR product of 105bp, 1ul of converted DNA and AmpliTaq Gold™ 360 Master Mix (Applied Biosystems, #4398876).

The PCR product was then run on 2% agarose (Sigma, #A4718) gel containing Ethidium Bromide (Sigma, #E1510) and picture were taken using Chemidoc XRS system (Biorad, # 1708265) with the single channel ethidium bromide agarose gel protocol. Band intensity was measured using Image J (https://imagej.nih.gov/ij/, 1997-2018. Schneider, C.A., Rasband, W.S., Eliceiri, K.W. “NIH Image to ImageJ: 25 years of image analysis”) and relative intensity between methylated and unmethylated specific PCR was calculated.

### Drugs and drug treatment

Cis-Diamineplatinium (II) dichloride (Cisplatin) (Sigma, #479306) was reconstituted fresh at 2mM in NaCl solution 0.9% (Sigma, #SW8776) and used at various concentration as indicated; Talazoparib (#S7048-Selleckchem) was reconstituted at 10mM in DMSO and used at various concentration.

### CRISPR cell engineering

#### Cloning of gRNAs into UNISAM plasmid

PB-UniSAM containing mCherry was a gift from Lesley Forrester (Addgene plasmid # 99866; http://n2t.net/addgene:99866; RRID: Addgene_99866) [19]. gRNAs were cloned into PB-UniSAM as previously published [18].

#### CRISPR cell engineering - Cell transfection

Electroporation was performed using Neon transfection system (Invitrogen, #MPK5000) with Neon™ Transfection System 10 µL Kit (Invitrogen, #MPK1096) using 2µg of DNA for 1.10^5^ cells in 24 well plate. Electroporation protocol for each cell line was identified using pmaxCloningTM vector (Lonza, # VDC-1040). The optimal protocol for each cell line is indicated in table 3 below:

**Table 3:** electroporation conditions for each cell line. After 1 week of culture, cells were further purified by sort.

| Cell line | Voltage | Width | Pulses |
| --- | --- | --- | --- |
| **PDM 182** | 1400 | 20 | 2 |
| **HTB 186** | 1150 | 30 | 2 |
| **CRL 1544** | 1700 | 20 | 1 |

### CRISPR cell engineering - Sort of cells

Cells were harvested and blocked in PBS with 5%FBS and 1%BSA and cell clumps removed on 40uM cell strainer (Falcon, #382235). Single-cell suspension was analyzed and sorted on SORP FACSAriaIII (BD Biosciences Special Order Research Product). Data were processed with BD FACSDiva™ Software V8.0.1 (BD Biosciences). mCherry fluorescence was acquired with 561 nm yellow-green laser and 610/20 nm emission filter. During cell-sorting 4-way purity-phase mask was applied. To ensure maximum purity, cells were serially sorted 3 times prior analysis and use.

### Viability analysis

Cells were grown in 96 well plate for 48 to 72 hours accordingly with or without treatment. 1500 cells per well were plated for PDM 182, HTB 186 and CRL 1544 respectively 24h prior treatment. Viability was assessed using ATPlite Luminescence Assay System (Perkinelmer, #6016949). Luminescence was measured with Ensight plate reader (Perkinelmer, #HH34000000).

### RNA seq analysis

#### mRNA sequencing

mRNA-sequencing was performed using QuantSeq 3’ mRNA-Seq Library Prep Kit FWD for Illumina (Cat. 015.96) (75 single-end) with a read depth of average 10.4 M, and average read alignment of 74%. Single samples were sequenced across multiple lanes, and the resulting FASTQ files were merged by sample. All samples passed FastQC (v.0.11.8) were aligned to the reference genome GRChg38 using STAR (v. 2.6.1d) [20]. BAM files were converted to a raw counts expression matrix using HTSeq-count (v.0.9.1) [21].

#### RNAseq Data Normalization

Normalization was done using R Bioconductor package EDAseq (Exploratory Data Analysis and Normalization for RNA-Seq) (v. 2.40.0) [22] to remove within and between lane effects. Data was then quantile normalized using R Bioconductor package preprocessCore package (v. 1.68.0) [23] and log2 transformed. All downstream analysis was done using R (v.4.4.1). Principal component analysis (PCA) was done based on genes expression to assess global transcriptional differences between the samples using prcomp() function and plotted using R CRAN package ggplot2 (v.3.5.2) [24].

#### Differentially expressed genes

Differentially expressed genes (DEGs) analysis was performed on log2 normalized mRNA expression data using R Bioconductor package limma (v.3.62.2) [25] with Benjamini-Hochberg (B-H) FDR. Within each comparative analysis, genes with row sum equal to zero were removed. To visualize the overlap of differentially expressed genes between the conditions, R CRAN package VennDiagram (v.1.7.3) was used [26]. Differentially expressed genes were then plotted in a heatmap using R Bioconductor package ComplexHeatmap (v.2.22.0) [27].

#### Pathway enrichment analysis

For enriched pathway analysis, list of DEGs (FDR < 0.01, LogFC >= 1) was uploaded to consensus path DB (CPDB) database or Ingenuity Pathways Analysis (IPA) to get the list of enriched pathways. For CPDB, output data was then downloaded as an ORA file and loaded to R studio for plotting bar plot using func2vis (v.1.0-3) package. For IPA, the list of pathways was exported as an excel file and used to regenerate the plot using ggplot2 (v. 3.5.2) package.

### Public data acquisition and analysis

#### Data acquisition and normalization

RNA-seq data for 4 pediatric tumors: Wilms tumor (WLM), neuroblastoma (NBL), osteosarcoma (OS) and rhabdoid tumor (RT) from the TARGET pediatric dataset, which is published on the GDC portal (https://portal.gdc.cancer.gov/), were downloaded and processed along with the clinical data as previously described [28]. The medulloblastoma RNAeq dataset was downloaded from the ICGC (https://dcc.icgc.org/).

#### Survival analysis

We performed survival analysis using survival (v. 2.41–3) and plotted the Kaplan-Meier curves using the ggsurv function from survminer (v. 0.4.8) [48]. For each cancer type we categorized the patient based on the log2 expression of SLFN11 into tertiles (High, Medium, Low). The overall survival analysis was conducted for each cancer type between the tertiles using the Cox proportional hazards regression analysis, as shown in Figure 1.

**Figure 1:**
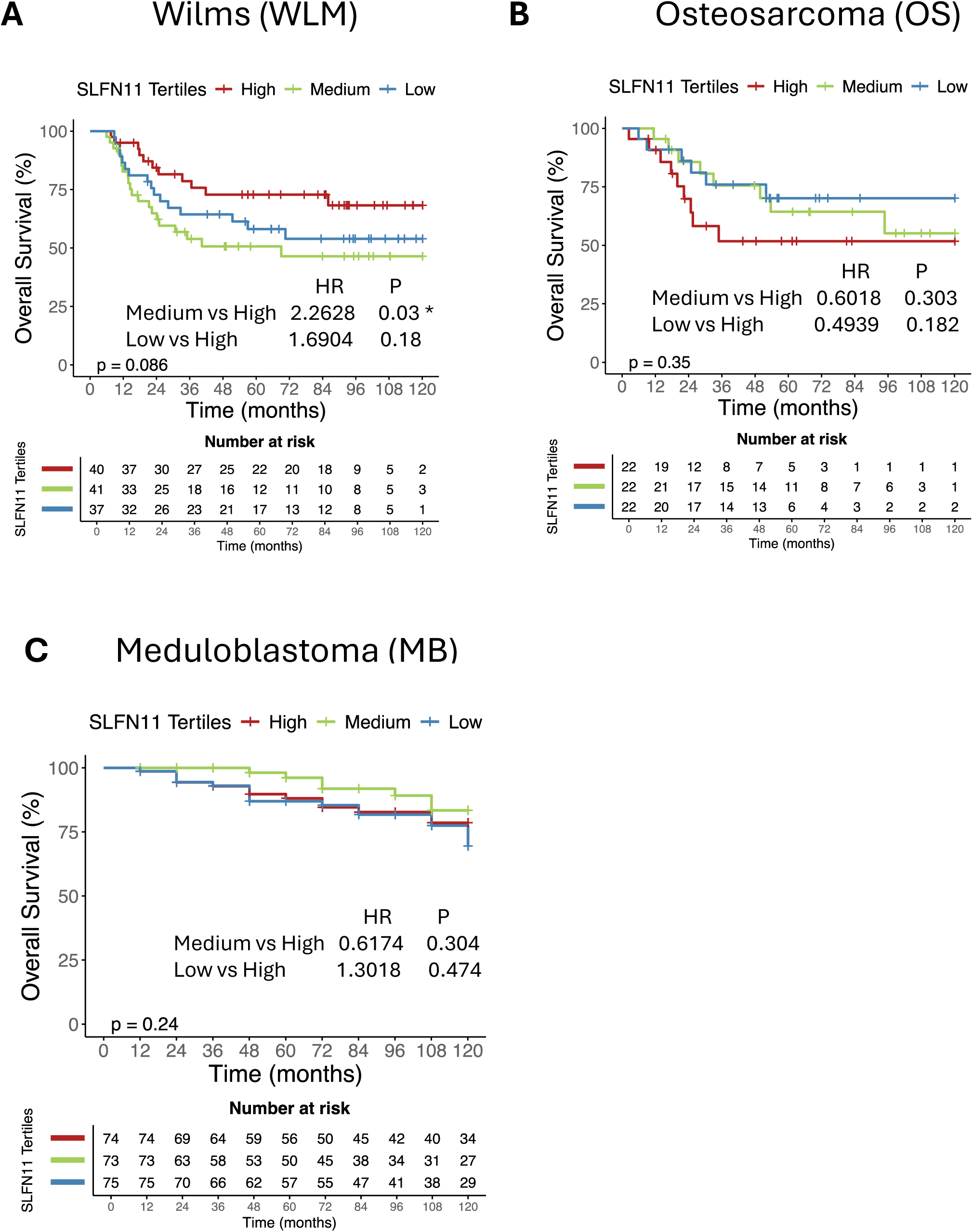
Tertials of SLFN1 expression in different pediatric solid tumors, and their prognosis from online TARGET and ICGC datasets. Kaplan-Meier survival curves showing overall survival stratified by SLFN11 expression tertiles (High, Medium, Low) in (A) Wilms Tumor (WLM), (B) Medulloblastoma (MB), and (C) Osteosarcoma (OS) from the TARGET pediatric cancer dataset. Hazard ratios (HR) and corresponding p-values for Medium vs High and Low vs High SLFN11 expression groups are indicated. (A) In Wilms Tumor, medium SLFN11 expression is associated with significantly worse overall survival compared to high expression (HR = 2.26, *p* = 0.03), (B) In Medulloblastoma, no significant association is observed between SLFN11 expression and survival (*p* = 0.35). (C) In Osteosarcoma, SLFN11 expression does not significantly affect overall survival (*p* = 0.24). The number of patients at risk at each time point is indicated below each graph.

### Statistical analysis

For statistical analysis and graphical presentation, Graphpad prism V10.1.0 (Domatics) software was used. Numerical results are given as means ± SD (N=sample size). The statistical significance for CWB and Q-PCR was assessed with Graphpad with unpaired Student’s t test or one way Anova. The statistical significance for the comparison of genes expression and enrichment score was calculated using unpaired t test using R programming function “stat_compare_means” from ggpubr package. Statistical significance was accepted for *p<0.05; **p<0.01; ***p<0.001; ****p<0.0001. Pearson correlation was performed using cor() R base function and plotted using corrplot() function from corrplot R package (v.0.92).

## Results

### Bioinformatic analysis of SLFN1 role in pediatric cancer

All types of pediatric cancers in the TARGET and ICGC dataset were analyzed *in-silico* to evaluate the predictive value of SLFN11. Patients were categorized into tertiles based on the log2 expression of SLFN11, and overall survival analysis was conducted for each cancer type, as shown in Figure 1. We selected three specific cancer types—Wilms Tumor, Osteosarcoma, and Medulloblastoma—because they exhibited different predictive outcomes. In Wilms Tumor (WLM), higher SLFN11 expression correlated with better survival rates as documented in many adult tumors. In contrast, Osteosarcoma (OS) showed better survival outcomes in patients with lower SLFN11 expression. The correlation in Medulloblastoma was neutral, showing no clear relationship (**Figure 1**). Based on these conflicting observations in the different tumor types, we decided to investigate, *in-vitro,* in these three-tumor types, the impact SLFN11 expression had on response to DNA damaging agents.

### Baseline SLFN1 expression and associated methylation profiles across a panel of 10 pediatric cancer cell lines

To establish baseline SLFN11 expression levels, we analyzed 9 pediatric cancer cell lines representing the selected cancer types. An immortalized human fibroblast cell line (HFF) was included as a normal control. SLFN11 expression was assessed at both mRNA and protein levels. Quantitative PCR (qPCR) analysis revealed differential SLFN11 mRNA expression across the cell lines. Among the two Wilms tumor cell lines screened, only PDM 182 exhibited detectable SLFN11 expression. All osteosarcoma cell lines showed some level of SLFN11 expression, while among the medulloblastoma cell lines, HTB186 demonstrated strong expression, and the remaining three displayed limited to no expression (**Figure 2A**).

**Figure 2:**
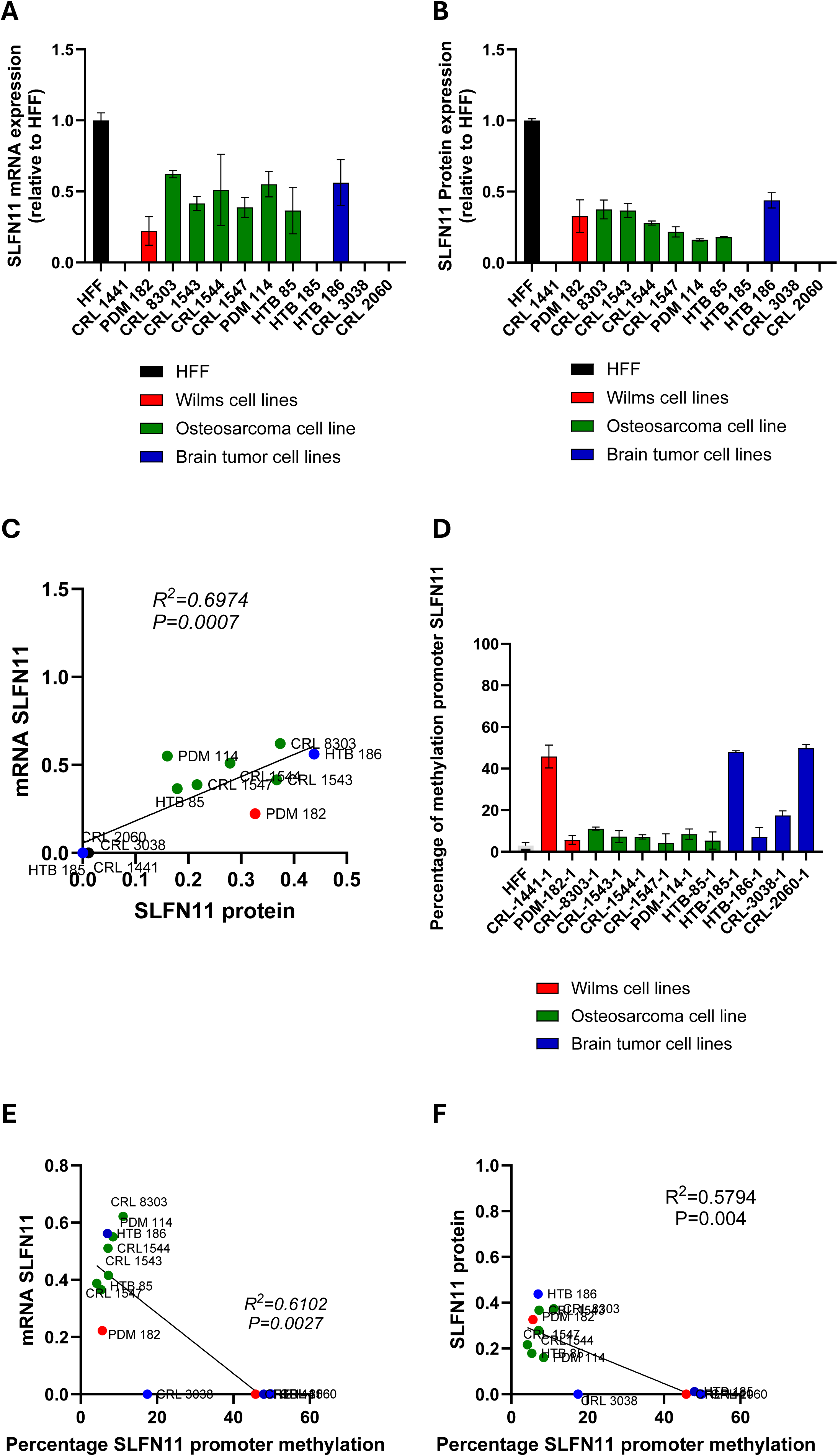
Baseline expression of SLFN1 across a panel of 9 pediatric cancer cell lines and associated methylation profile. (A) Relative mRNA expression of SLFN11 analyzed by Q-RT-PCR. SLFN11 expression in the different cell lines shown relative to the expression level in HFF (human foreskin fibroblast) used as control cells (N=3). (B) Capillary Western blot immunoassay (CWB) of SLFN11 expression in those breast cancer cell lines. Quantification of band intensity of SLFN11 relative to the expression level in HFF cells (N=2). (C) Correlation between SLFN11 mRNA analyzed by Q-RT-PCR and protein expression analyzed by CWB. (D) Percentage of methylation of SLFN11 promoter analyzed by methyl specific PCR (MSP) within the CPG island of the promoter (N=3). (E). Correlation of SLFN11 promoter methylation and SLFN11 mRNA expression analyzed by Q-RT-PCR. (F). Correlation of SLFN11 promoter methylation and SLFN11 protein expression analyzed by CWB.

To validate these findings at the protein level, we performed CWB analysis (**Figure 2B**) (**Supplementary Figure 1**). The protein expression patterns was significantly correlated with the mRNA expression levels observed in qPCR. A significant correlation between SLFN11 mRNA and protein expression was confirmed (R²=0.70, p=0.0007) (**Figure 2C**), indicating consistent transcriptional and translational regulation.

Given that promoter methylation is a key regulatory mechanism of SLFN11 expression, we next analyzed SLFN11 promoter methylation using methylation-specific PCR (MSP) (**Figure 2D**). Significant correlations were observed between promoter methylation and both mRNA (R²=0.61, p=0.0027) and protein (R²=0.58, p=0.004) expression levels (**Figure 2E and 2F**). These findings suggest that promoter methylation plays a critical role in regulating SLFN11 expression in pediatric cancer cell lines.

The strong correlation between SLFN11 expression and promoter methylation aligns with previous findings in adult cancer cell lines, including our observations in breast cancer models, with no expression when strong methylation is observed. This indicates that SLFN11 expression in pediatric cancers may be regulated by mechanisms similar to those in adult cancers, highlighting its potential as a conserved biomarker across cancer types.

### CRISPR-dCas9 effectively modulates SLFN1 expression in selected pediatric cancer cell lines

To modulate SLFN11 expression, we employed the UNISAM (Unique Synergistic Activation Mediator) system using dead Cas9 (dCas9) activation, a method we previously validated in breast cancer cell lines [18,19]. Based on our earlier findings, we selected one cell line from each cancer type (Wilms tumor, osteosarcoma, and medulloblastoma) with minimal but detectable SLFN11 expression. This selection ensured that CRISPR activation could effectively increase SLFN11 levels without being hindered by strong promoter methylation, which we previously found to be a limiting factor that UNISAM could not overcome [18]. The cell lines chosen are PDM-182 (Wilms tumor), CRL-1544 (osteosarcoma), and HTB-186 (medulloblastoma).

We utilized the same gRNAs previously designed to target the central region of the SLFN11 promoter (**Table 2**) [18]. Stable cell lines expressing five SLFN11-targeting gRNAs and a scrambled (SCR) control gRNA were generated. Following selection and purification based on mCherry expression (a UNISAM reporter), all modified cell lines were screened using CWB and qPCR to identify the most effective gRNA for modulating SLFN11 expression.

In PDM-182, gRNA N4 emerged as the most efficient, as confirmed by both qPCR and CWB (**Supplementary Figure 2 A-B**). For CRL-1544 and HTB-186, gRNA N10 was identified as the most effective (**Supplementary Figure 2 C-F**). Further validation was performed using CWB (N=6; 3 biological replicates, 2 technical replicates each) and qPCR (N=9; 3 biological replicates, 3 technical replicates each). A significant increase in SLFN11 mRNA and protein levels was observed with the SLFN11 specific gRNAs (P<0.0001 in all cell lines), while no significant change was detected with the scrambled (SCR) gRNA (**Figure 3 A-F**).

**Figure 3:**
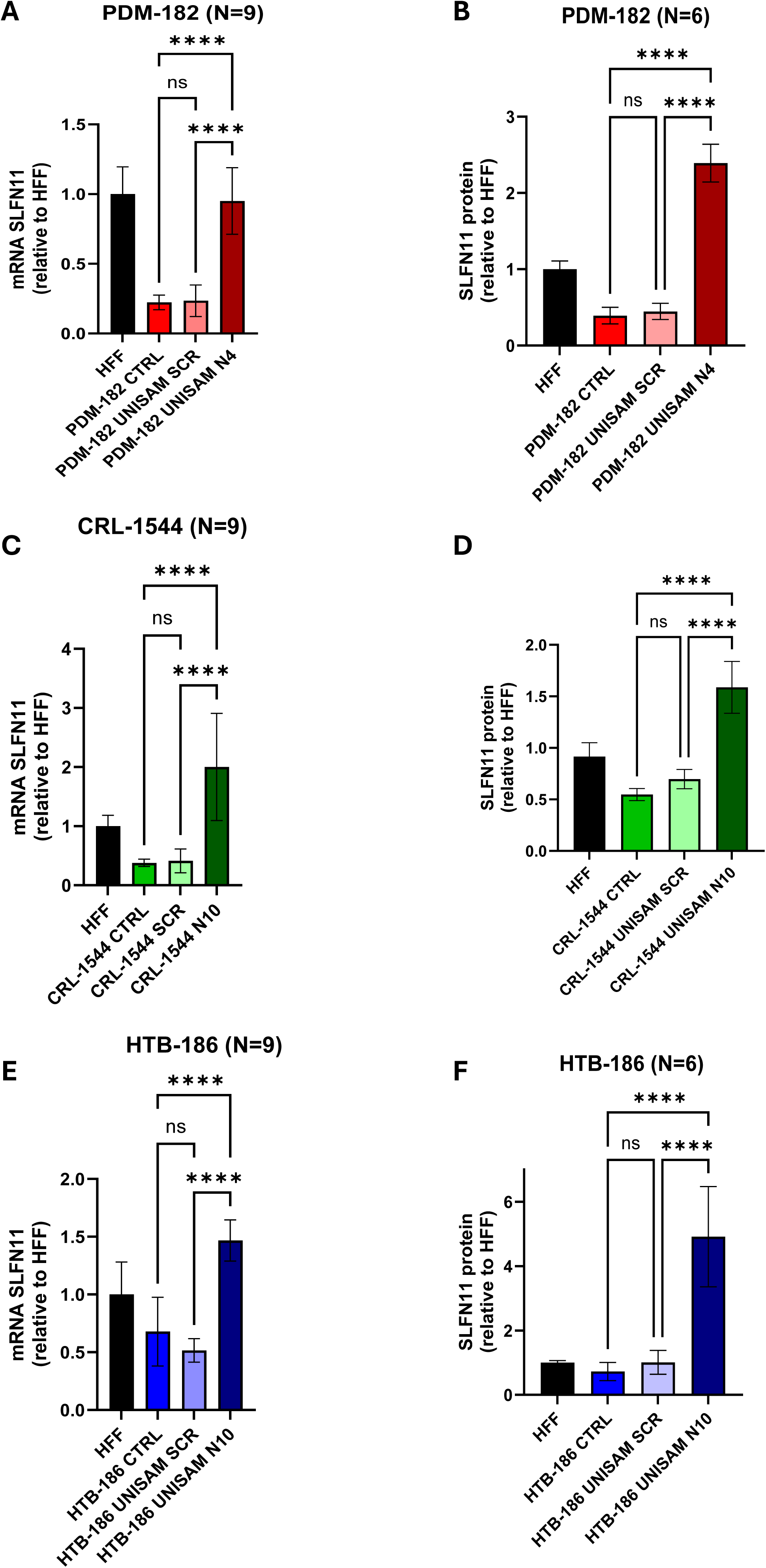
CRISPR-cas9 modulation of SLFN1 expression in 3 cell lines. The UNISAM (unique Synergistic Activation Mediator) system developed by Fidanza, A., et al. was used for CRISPR activation of SLFN11 (14). Using the gRNA N4 and gRNA N10 we could successfully increase SLFN11 expression in PDM182 as analyzed by Q-RT-PCR (N=9, 3 technical replicates of 3 biological replicates) (A) and CWB (N=6, 2 technical replicates of 3 biological replicates) (B). Similarly, Using the gRNA N7 and gRNA N10 we could successfully increase SLFN11 expression in CRL1544 as analyzed by Q-RT-PCR (N=9, 3 technical replicates of 3 biological replicates) (C) and CWB (N=6, 2 technical replicates of 3 biological replicates) (D). Finally, using gRNA N7 and gRNA N10 we could successfully increase SLFN11 expression in PDM186 as analyzed by Q-RT-PCR (N=9, 3 technical replicates of 3 biological replicates) (E) and CWB (N=6, 2 technical replicates of 3 biological replicates) (F). In opposition, the UNISAM and KRAB system used with scramble gRNA (SCR) did not significantly affect SLFN11 expression compared to respective untreated cells (CTRL).

These results demonstrate that UNISAM activation can effectively upregulate SLFN11 expression in pediatric cancer cell lines. Having established this, we next sought to determine whether increased SLFN11 expression enhances sensitivity to DNA-damaging agents (DDAs) such as cisplatin and a PARP inhibitor such as talazoparib.

### Modulation of SLFN1 enhances sensitivity to cisplatin and talazoparib treatment

To evaluate the impact of SLFN11 expression on chemosensitivity, CRISPR-modified cells and their control counterparts were treated with two DNA-damaging agents: cisplatin, a platinum-based chemotherapeutic, and a PARP inhibitor: talazoparib. Dose-response curves were generated for each drug, with treatment durations and concentration ranges tailored to the specific agent and cell line. Statistical significance was assessed at the concentration where near-maximal effects were observed in the most sensitive cell line, prior to reaching a toxicity plateau (Figure 4 B, D, F, H, J, L).

**Figure 4:**
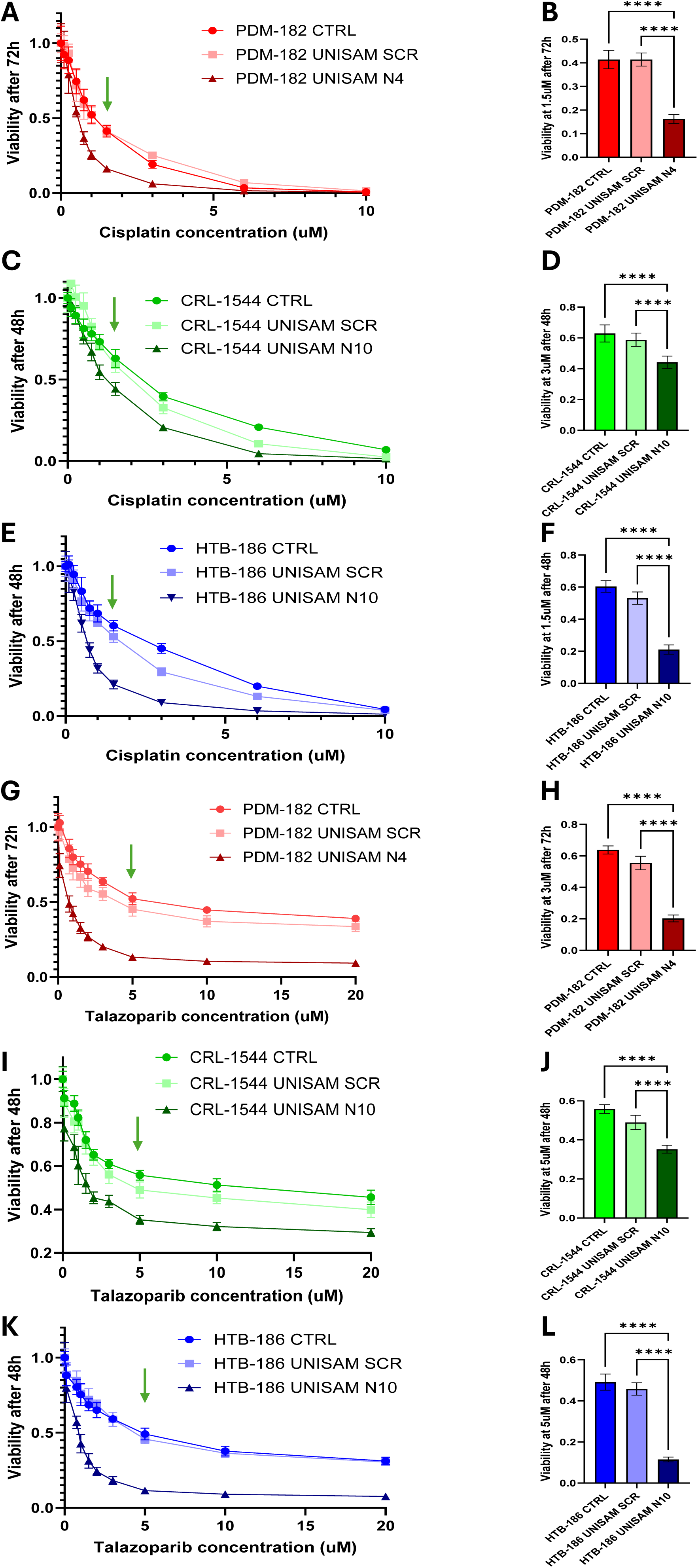
CRISPR-dCas9 modulation of SLFN1 impact sensitivity to DNA Damaging agents. SLFN11 increase with UNISAM (gRNA N10 for PDM-182 and HTB-186 and gRNA N7 for CRL-1544) leads to significant increase sensitivity to Cisplatin treatment (A,B for PDM-182, E,F for CRL-1544 and I,J for HTB-186) (N=18, 3 biological replicates of 6 technical replicates) and Talazoparib treatment (C,D for PDM-182, G,H for CRL-1544 and K,L for HTB-186) (N=18, 3 biological replicates of 6 technical replicates) compared to scramble (SCR) control and CTRL (unmodified cells).

#### Cisplatin treatment

PDM-182 cells (Wilms tumor) were treated with cisplatin at concentrations ranging from 0.1 µM to 20 µM for 72 hours. Cell viability was measured using the ATP Lite assay (N=18; 3 biological replicates, 6 technical replicates) (**Figure 4A**). Compared to untreated cells and cells modified with scrambled gRNA (UNISAM SCR), PDM-182 cells expressing the optimal gRNA (UNISAM N4) exhibited significantly increased sensitivity to cisplatin across a broad concentration range. At 1.5 µM, UNISAM N4-modified cells showed markedly higher sensitivity than UNISAM scrambled (SCR) and unmodified control (CTRL) cells (p<0.0001) (**Figure 4B**).

A similar trend was observed in CRL-1544 (osteosarcoma) cells treated with cisplatin for 48 hours. UNISAM N10-modified cells demonstrated enhanced sensitivity compared to UNISAM SCR and CTRL cells across multiple concentrations (**Figure 4C**). At 1.5 µM, CRL-1544 UNISAM N10 cells were significantly more sensitive than their scrambled and control counterparts (p<0.0001) (**Figure 4D**).

In HTB-186 (medulloblastoma) cells, cisplatin treatment for 48 hours also revealed increased sensitivity in UNISAM N10-modified cells (**Figure 4E**). At 1.5 µM, HTB-186 UNISAM N10 cells showed significantly greater sensitivity compared to UNISAM SCR and CTRL cells (p<0.0001) (**Figure 4F**).

It is noteworthy to point that to reach similar level of cell death 72h were necessary with PDM-182, while CRL-1544 and HTB-186 reached it in 48h.

#### Talazoparib treatment

Talazoparib, a potent PARP inhibitor, was used to treat PDM-182, CRL-1544, and HTB-186 cells at concentrations ranging from 0.1 µM to 20 µM. Treatment durations were 72 hours for PDM-182 and 48 hours for CRL-1544 and HTB-186. Consistent with the results on cisplatin, SLFN11-activated cell lines exhibited increased sensitivity to talazoparib compared to scrambled and control cells across various concentrations (**Figure 4G, I, K**). For example, at 5 µM talazoparib for 72 hours, PDM-182 UNISAM N4 cells demonstrated significantly greater sensitivity than UNISAM SCR and CTRL cells (p<0.0001) (**Figure 4H**). Similarly, at 5 µM talazoparib for 48 hours, both CRL-1544 UNISAM N10 and HTB-186 UNISAM N10 cells showed significantly increased sensitivity compared to their scrambled and control counterparts (p<0.0001) (**Figure 4J, L**).

#### RNAseq analysis of CRISPR modified cells and treatment with cisplatin

Principal component analysis (PCA) revealed that UNISAM-SCR and UNISAM-N (targeting SLFN11 with different gRNA depending on cell lines as presented before) samples clustered closely together under untreated conditions across all three cell lines, indicating minimal basal transcriptional differences (**Figure 5A**). Under cisplatin treatment, cells PCA shift (represented by the arrows for each cell line) from untreated, is captured by a different dimension in the PCA plot.

**Figure 5:**
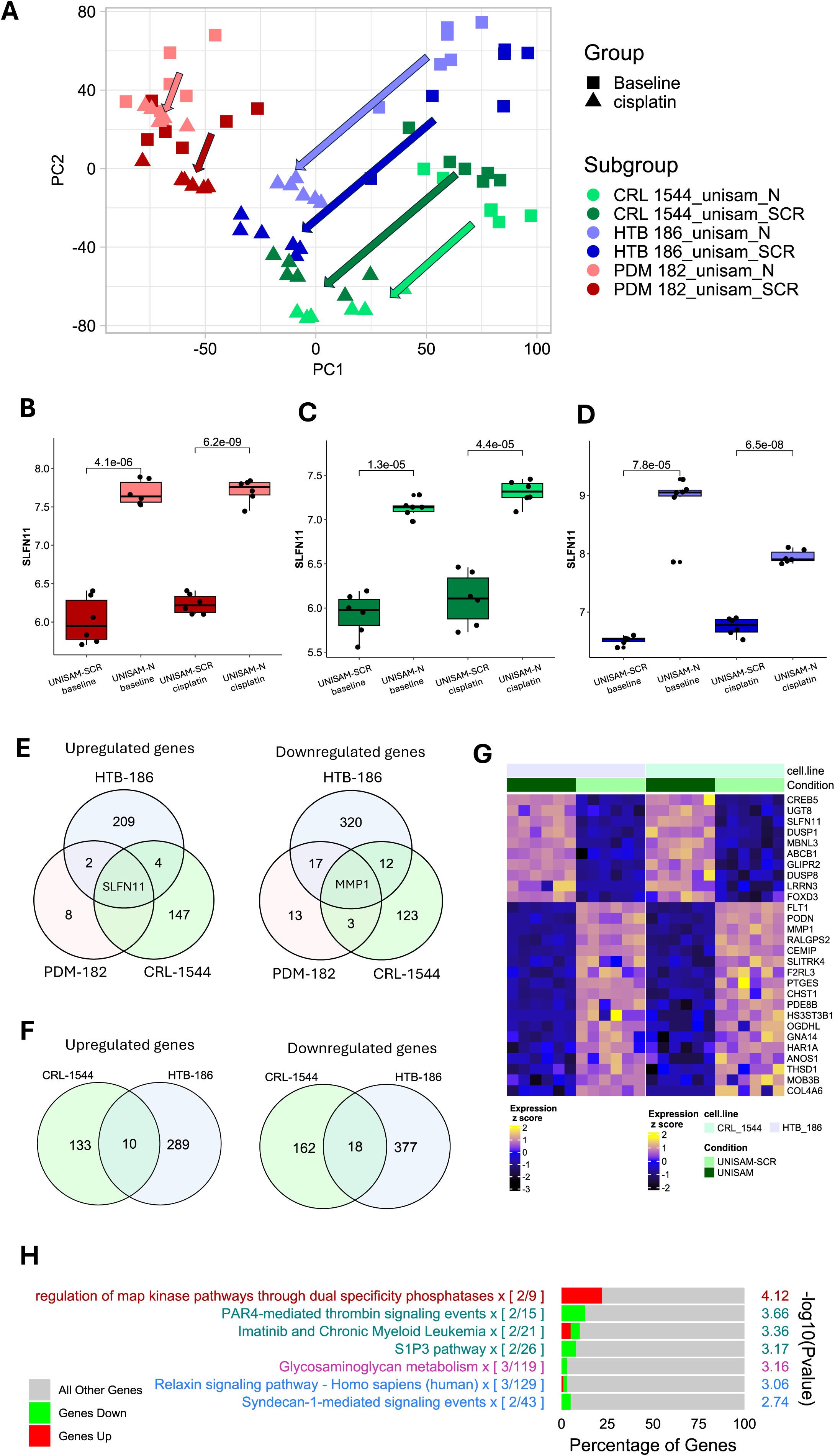
RNAseq analysis of CRISPR modified cells and cisplatin treatment. (A). Principal component analysis based on gene expression for all samples (baseline and cisplatin-treated samples) PDM-182 (red), CRL-1544 (green) and HTB-186 (blue) cell lines. (B, C, D). Expression of SLFN11 across baseline and cisplatin-treated samples in PDM-182, CRL-1544 and HTB-186 cell lines respectively. Statistical significance was assessed using an unpaired t test. (E). VennDiagram of common differentially expressed genes between all three cell lines which are upregulated (N = 1, SLFN11) and downregulated (N = 1, MMP1) between UNISAM-SCR and UNISAM-N (targetin SLFN11) without treatment ((FDR < 0.01, logFC < |1|)). (F). VennDiagram of common differentially expressed genes between CRL-1544 and HBT-186 cell lines which are upregulated (N = 10) and downregulated (N = 18) between UNISAM-SCR and UNISAM-N (targeting SLFN11) under cisplatin treatment ((FDR < 0.01, logFC < |1|)). (G) Heatmap of those common differentially expressed genes. (H) ConsensusPathDB analysis of the 28 genes previously identified.

In alignment with our Q-RT-PCR and western blot analysis, RNA-seq data further confirmed that SLFN11 expression is consistently upregulated in UNISAM-N compared to UNISAM-SCR across all three cell lines, both in the presence and absence of cisplatin (**Figure 5B-D**). Importantly, SLFN11 was the only gene upregulated across all three untreated UNISAM-N cell lines, highlighting the specificity of the UNISAM construct in activating SLFN11 (**Figure 5E**). Interestingly, we observed MMP1 was also systematically downregulated across all three cell lines.

Differential expression analysis between untreated and cisplatin-treated conditions confirmed a robust cisplatin-induced transcriptional response in each cell line, characterized by enrichment of gene sets related to DNA replication, mismatch repair, and DNA damage response pathways, as determined by the consensus pathway database (CPDB) analysis (**Supplementary figure 3**). Yet under cisplatin treatment, comparison of UNISAM-SCR vs. UNISAM-N revealed varying numbers of differentially expressed genes (DEGs) across the cell lines. Similar to the PCA, PDM-182 showed minimal changes, while HTB-186 and CRL-1544 exhibited substantial transcriptomic differences. This aligns with previous findings: PDM-182 requires a 72-hour cisplatin treatment to observe a notable toxicity differential effect between UNISAM-SCR and UNISAM-N, whereas HTB-186 and CRL-1544 display significant cell death as early as 48 hours. These results suggest that PDM-182 is less sensitive to cisplatin, and therefore, 24-hour treatment may be insufficient to capture the SLFN11-mediated transcriptional effects in this line. This reduced cisplatin sensitivity in PDM-182 is further illustrated by the minimal PCA shift observed under treatment compared to the more prominent shifts in HTB-186 and CRL-1544 (**Figure 5A**). As a result, subsequent analysis focused on HTB-186 and CRL-1544 only.

Upon cisplatin treatment, both UNISAM-SCR and UNISAM-N samples in both cell lines (HTB-186 and CRL-1544) showed a pronounced shift in the same direction, suggesting a common transcriptomic response to DNA damage (**Supplementary figure 4A-D**)).

When comparing UNISAM-SCR vs. UNISAM-N under cisplatin treatment in HTB-186 and CRL-1544, we identified 10 genes upregulated and 18 genes downregulated in common between the two cell lines (**Figure 5F**). This list is highlighted in the heatmap (Figure 5G). With this gene list we performed a CPDB analysis and identified as significantly upregulated MAPK pathways activation, while PAR-4-mediated thrombin signaling events signaling pathway and S1P3 pathway as downregulated (**Figure 5H**). Similarly, Ingenuity pathway analysis (IPA) showed in the most significant pathways RAF-independent MAPK1/3 pathway activation and p38 MAPK signaling upregulation while thrombin signaling (PAR), dermatan/chondroitin/heparan sulfatan biosynthesis and collagen degradation were down regulated (**Supplementary figure 5**). In the UNISAM-N dataset, analysis of MAPK pathway regulator gene expression in HTB-186 and CRL-1544 cells without cisplatin treatment reveals a significant negative correlation with DUSP8 (r= −0.76, p=0.0044) and MAPK3 (r= −0.9, p=5.7e-5) (**Supplementary figure 6)**. The same dataset under cisplatin treatment revealed a positive correlation (r ≥ 0.7) between SLFN11 and the phosphatases DUSP1 (0.002), DUSP4 (p=4.1e-5), and DUSP8 (p=0.0018), and a negatively correlated (r ≤ −0.7) with DUSP6 (0.002) and MAPK14 (p=0.000441), indicating potential pathway divergence or inhibitory crosstalk between SLFN11-mediated DNA damage response and MAPK signaling components such as p38 MAPK. (**Supplementary figure 7**).

## Discussion

Pediatric cancers remain a therapeutic challenge due to their biological heterogeneity and sometimes limited responsiveness to standard chemotherapeutics. Among these, Diffuse Intrinsic Pontine Glioma (DIPG) stands out as one of the most aggressive and least survivable pediatric cancers. DIPG is a rare brain tumor that primarily affects children and has a median survival of less than one year, with no effective cure currently available [29]. Wilms tumor is a treatable pediatric kidney cancer with a 5-year survival rate exceeding 90%, especially in patients with favorable histology [30]. On the other end, Osteosarcoma and medulloblastoma have more variable outcomes; osteosarcoma has about a 65% 10-year survival, while medulloblastoma survival ranges widely (up to 100% in low-risk subtypes, but significantly lower in high-risk cases), depending on molecular subtype and treatment response [31,32]. While significant progress has been made in adult oncology in identifying biomarkers of chemosensitivity, such precision-guided strategies are still underdeveloped in pediatric settings. In this study, we demonstrate that SLFN11, a DNA/RNA helicase-like protein previously linked to enhanced sensitivity to DNA-damaging agents in adult tumors [1,33] plays a functionally conserved yet context-dependent role in pediatric tumors. Through integrated in silico, transcriptional, and functional validation approaches, we provide evidence that SLFN11 expression can serve as a predictive biomarker for response to genotoxic chemotherapy in specific pediatric malignancies, and that its epigenetic silencing may contribute to treatment resistance [34].

Our work expands on prior adult cancer studies [18] by examining SLFN11 in the context of pediatric medulloblastoma, osteosarcoma, and Wilms tumor—three tumor types that reflect divergent SLFN11 expression profiles and treatment responses observed in silico. We established that upregulation of SLFN11 expression increase chemosensitivity to platinum-based agents and PARP inhibitors [35,36]. These findings offer a compelling rationale for reactivating SLFN11 in pediatric tumors that exhibit intrinsic or acquired chemoresistance. Importantly, although SLFN11 is not currently recognized as a predictive marker of chemosensitivity in pediatric tumors, its functional role appears to reflect that observed in adult tumors.

We investigated the molecular mechanisms by which SLFN11 enhances cellular sensitivity to cisplatin, a DNA-damaging chemotherapeutic using RNA sequencing. SLFN11-overexpressing cells displayed significantly increased sensitivity to cisplatin. To delineate the signaling pathways involved, we performed transcriptomic profiling followed by pathway enrichment analysis using both ConsensusPathDB and Ingenuity Pathway Analysis (IPA). The analyses revealed enhanced MAPK signaling and a reprogramming of the cellular stress response to favor apoptosis over senescence. Additionally, SLFN11 expressions was associated with suppression of pro-survival and chemoresistance-associated pathways, including PI3K/AKT and NF-κB signaling. These findings suggest that SLFN11 sensitizes cells to cisplatin by shifting the stress-response landscape to promote apoptotic cell death and diminish resistance mechanisms.

Pathway analysis highlighted robust activation of several MAPK-related cascades, including RAF-independent MAPK1/3 (ERK1/2) activation, p38 MAPK signaling, and the ERK/MAPK pathway, which are key regulators of the DNA damage response, cell cycle arrest, and apoptosis. Interestingly, without cisplatin, SLFN11 expression correlates positively with DUSP4 and negatively with MAPK3, suggesting that SLFN11 helps regulate MAPK signaling by promoting phosphatase activity and suppressing proliferative kinase expression. Under cisplatin treatment, SLFN11 shows positive correlations with multiple other phosphatases (DUSP1, DUSP4, DUSP8) and negative correlations with DUSP6 and MAPK14 (p38), indicating a selective modulation of MAPK components. This pattern suggests that SLFN11 may enhance phosphatase-mediated inhibition of MAPK signaling while limiting p38 MAPK activation, thus fine-tuning the DNA damage response through inhibitory crosstalk with specific MAPK pathways. While DUSPs can deactivate MAPKs, their selective expression can paradoxically sensitize cells to stress by shaping ERK and p38 dynamics to favor apoptosis, as previously described [37,38].

Simultaneously, pro-survival and chemoresistance-associated pathways were downregulated. Notably, both analyses identified a suppression of thrombin signaling through protease-activated receptors (PARs), particularly PAR-4, which is involved in thrombin-induced inflammatory and survival signaling. Inhibition of this pathway may diminish tumor-promoting inflammation and cellular resistance to genotoxic stress [39]. Furthermore, the S1P3 signaling pathway, which typically activates PI3K/AKT and ERK to promote survival, migration, and chemoresistance, was also repressed, pointing to a broader suppression of pro-survival signaling [40–42].

Additional pathways downregulated in SLFN11-overexpressing cells under cisplatin treatment included chondroitin sulfate and heparan sulfate biosynthesis, which are key components of the extracellular matrix and known to modulate tumor progression, metastasis, and drug accessibility [43,44]. Their inhibition may contribute to a less protective tumor microenvironment. Finally, changes in drug metabolism pathways, such as atorvastatin and prednisone ADME (Absorption, Distribution, Metabolism, and Excretion), were noted, suggesting broader alterations in cellular homeostasis and response to xenobiotics, potentially enhancing cisplatin accumulation or efficacy [45–47]. Together, these findings indicate that SLFN11 expression in those pediatric cancer enhances cisplatin sensitivity through a coordinated rewiring of intracellular signaling, involving activation of stress-related MAPK pathways (including ERK1/2 and p38), and suppression of multiple pro-survival and extracellular resistance mechanisms. This dual action creates a cellular landscape more prone to apoptosis and less capable of counteracting DNA damage, providing a mechanistic rationale for the use of SLFN11 as a predictive biomarker and potential therapeutic modulator in cisplatin-based chemotherapy.

Without cisplatin treatment, the observed downregulation of MMP1 when SLFN11 is upregulated suggests that SLFN11 may act to suppress genes involved in extracellular matrix remodeling and invasion. Since MMP1 is a matrix metalloproteinase that promotes tissue degradation and tumor progression, its repression by SLFN11 could reflect a tumor-suppressive role, helping to limit invasive potential in cells with higher SLFN11 expression under basal conditions.

## Conclusion

Our integrated bioinformatic and experimental study establishes SLFN11 as a key modulator of chemosensitivity in pediatric solid tumors, with tumor type–specific predictive value. While high SLFN11 expression correlates with improved survival and enhanced sensitivity to DNA-damaging agents in Wilms tumor, its role seems more complex in osteosarcoma and medulloblastoma, reflecting an apparent tumor-specific biology. We demonstrate that epigenetic regulation via promoter methylation critically controls SLFN11 expression across these pediatric cancer type cell lines, and that CRISPR-dCas9–mediated activation effectively increases SLFN11 levels, thereby increasing sensitivity to cisplatin and PARP inhibition in all tested cancer types. Transcriptomic analyses suggest SLFN11’s involvement in modulating DNA damage response and MAPK signaling pathways, highlighting potential mechanisms underlying its chemo sensitizing effect. These findings underscore SLFN11’s promise as a biomarker and therapeutic target for enhancing response to genotoxic chemotherapy in pediatric oncology, warranting further clinical investigation and development of SLFN11-targeted strategies to overcome chemoresistance in childhood cancer.

## List of abbreviations

DDAs: DNA damaging agents
DDR: DNA Damage Repair
ADP: Adenosine diphosphate-ribose
PARPi: Poly ADP polymerase inhibitor
RPA: Replication Protein A
MCM3: Minichromosomal maintenance complex component 3
DHX9: DExH-box helicase
IFN-γ: interferon-gamma
SCLC: small cell lung cancer

## Acknowledgements

The authors would like to acknowledge the Sidra Medicine research branch core facilities and our research administration team without whom we could not have performed this work.

## Declarations

### Ethics approval and consent to participate

“Not applicable”

### Consent for publication

“Not applicable”

### Author’s contribution

C.R., D.A., A.J., performed the experiments, E.A., S.S., R.A., performed the computational analysis C.R., drafted the manuscript and the figures, W.H., C.R, E.S. edited the text. Conceptualization by S.S., C.R. and W.H., project supervision and coordination C.R., W.H. All authors reviewed the manuscript.

## Figure caption

*Supplementary figure 1*

Representative CWB results of the analysis of SLFN11 protein expression in the 11 tested unmodified pediatric cancer cell lines compared to human foreskin fibroblast (HFF).

*Supplementary figure 2*

(A,C, E) Relative mRNA expression of SLFN11 analyzed by Q-RT-PCR (n=3) (B,D,F) and relative SLFN11 protein expression analyzed by CWB (n=2) (B-D) in PDM-182 (A,B), CRL-1544 (C,D) and HTB-186 (E,F) cancer cell lines modified with each gRNA for CRISPR-dCas9-UNISAM relative to HFF.

*Supplementary figure 3*

ConsensusPathDB analysis of up and downregulated differentially expressed genes (FDR < 0.01, logFC < |1|) between untreated and treated with 1.5uM cisplatin UNISAM-SCR for each cell line (A: PDM-182, B: CRL-1544, C: HTB-186) showing a clear response to cisplatin treatment in SCR cells for CRL-1544 and HTB-186 with cell cycle arrest and DNA damage response, but not in PDM-182. Full list of obtained pathways were filtered to include only those with FDR < 0.01.

*Supplementary figure 4*

(A). VennDiagram of differentially expressed genes (FDR < 0.01, logFC < |1|) between untreated and 1.5uM cisplatin for 24h treated cells in the UNISAM-SCR of CRL-1544 and HTB-186 showing the genes triggered by cisplatin treatment without upregulation of SLFN11 (B) ConsensusPathDB analysis of those common up and downregulated differentially expressed genes for UNISAM-SCR showing a clear response to cisplatin treatment with cell cycle arrest and DNA damage response. Full list of obtained pathways were filtered to include only those with FDR < 0.01.

(C). VennDiagram of differentially expressed genes (FDR < 0.01, logFC < |1|) between untreated and 1.5uM cisplatin for 24h treated cells in the UNISAM-N (targeting SLFN11) of CRL-1544 and HTB-186 showing the genes triggered by cisplatin treatment without upregulation of SLFN11 (D) ConsensusPathDB analysis of those common up and downregulated differentially expressed genes for UNISAM-N also showing a clear response to cisplatin treatment with cell cycle arrest and DNA damage response. Full list of obtained pathways were filtered to include only those with FDR < 0.01.

*Supplementary figure 5*

Ingenuity Pathway Analysis of the 28 commonly differentially expressed genes between UNISAM-N and UNISAM-SCR (targeting SLFN11) under cisplatin treatment in both CRL-1544 and HTB-186 cell lines ((FDR < 0.01, logFC < |1|)). Histograms represent the proportion (%) of DEGs upregulated (red) or downregulated (green) in UNISAM-N vs. UNISAM-SCR. The circles represent the pathway activation status. Blue circle indicates the pathway is inhibited with a negative z-score, the white circle represents the pathway is neutral with zero z-score, while a gray circle indicates that the pathway activity is unknown.

*Supplementary figure 6*

(A) Pearson correlation plot between SLFN11 expression and proteins involved in regulation of map kinase pathway and DUSPs under 1.5uM cisplatin treatment after 24h in UNISAM-N of PDM-186 and CRL-1544. Shown values represent the correlation coefficients and p-values between the brackets. For the exact p-value of the non-significant (ns) please refer to the supplementary file 1, table 1.

*Supplementary figure 7*

(A) Heatmap of expression of proteins involved in regulation of map kinase pathway by DUSPs in PDM-186 and CRL-1544 of both UNISAM-SCR and UNISAM-N under 1.5uM cisplatin treatment after 24h (B) Pearson correlation plot between SLFN11 expression and proteins involved in regulation of map kinase pathway and DUSPs under 1.5uM cisplatin treatment after 24h in UNISAM-N of PDM-186 and CRL-1544. Shown values represent the correlation coefficients and p-values between the brackets. For the exact p-value of the non-significant (ns) please refer to the supplementary file 1, table 2.

