## Supplementary figures for "SLFN11 Enhances Cisplatin Sensitivity in Pediatric Cancer via Activation of Stress-Response and Suppression of Survival Pathways"

Supplementary figure 1

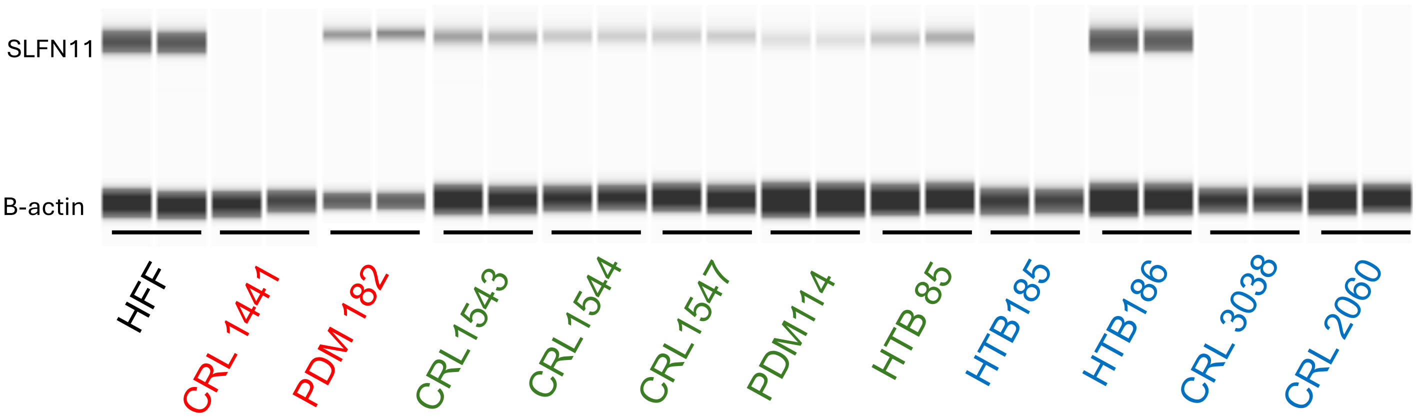

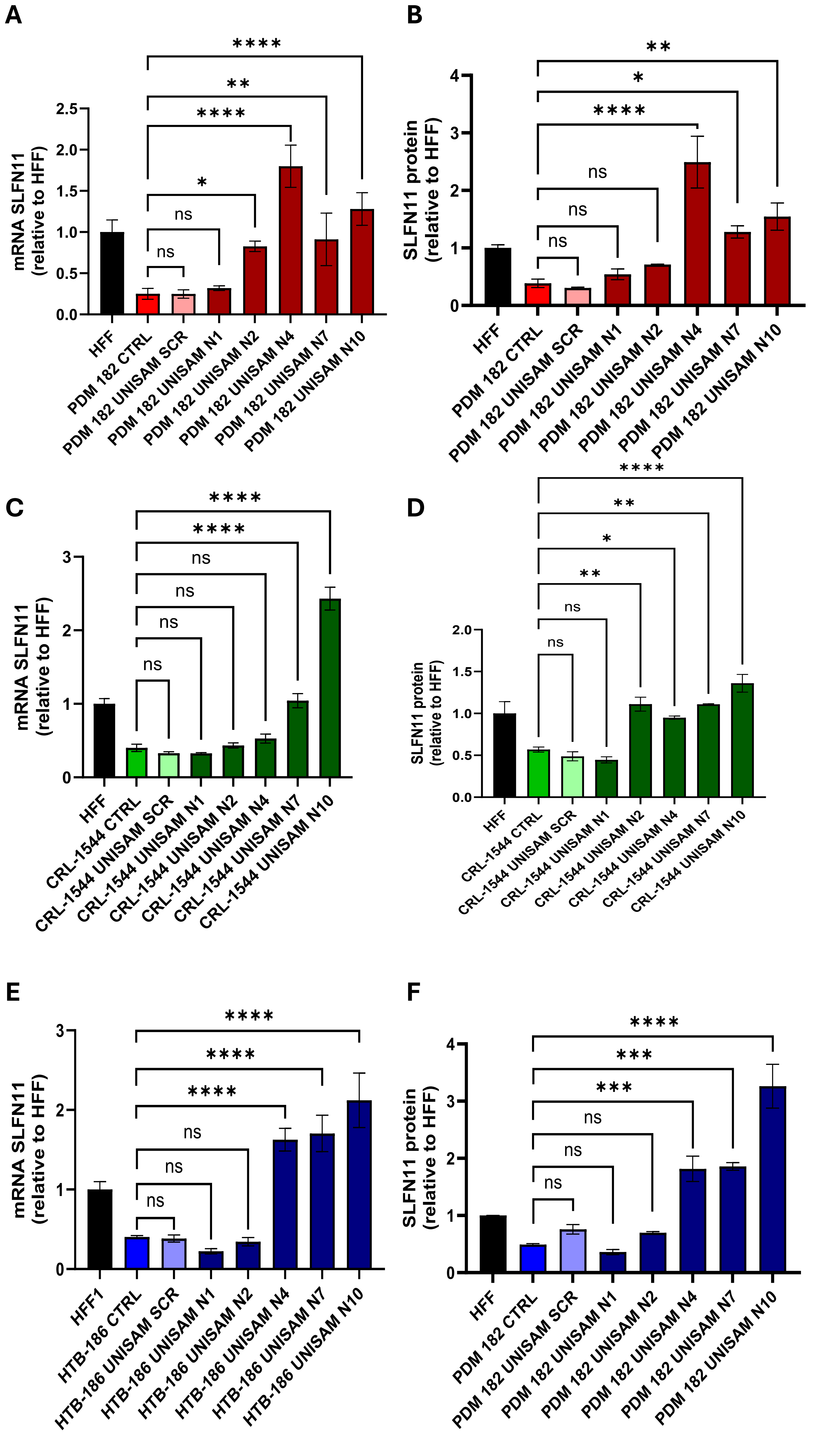

A
B
C
D
E
F
mRNA SLFN11 (relative to HFF)
SLFN11 protein (relative to HFF)
ns
*
**
***
****
HFF
HFF1
PDM 182 CTRL
PDM 182 UNISAM SCR
PDM 182 UNISAM N1
PDM 182 UNISAM N2
PDM 182 UNISAM N4
PDM 182 UNISAM N7
PDM 182 UNISAM N10
CRL-1544 CTRL
CRL-1544 UNISAM SCR
CRL-1544 UNISAM N1
CRL-1544 UNISAM N2
CRL-1544 UNISAM N4
CRL-1544 UNISAM N7
CRL-1544 UNISAM N10
HTB-186 CTRL
HTB-186 UNISAM SCR
HTB-186 UNISAM N1
HTB-186 UNISAM N2
HTB-186 UNISAM N4
HTB-186 UNISAM N7
HTB-186 UNISAM N10

### UNISAM vs. UNISAM-SCR Baseline

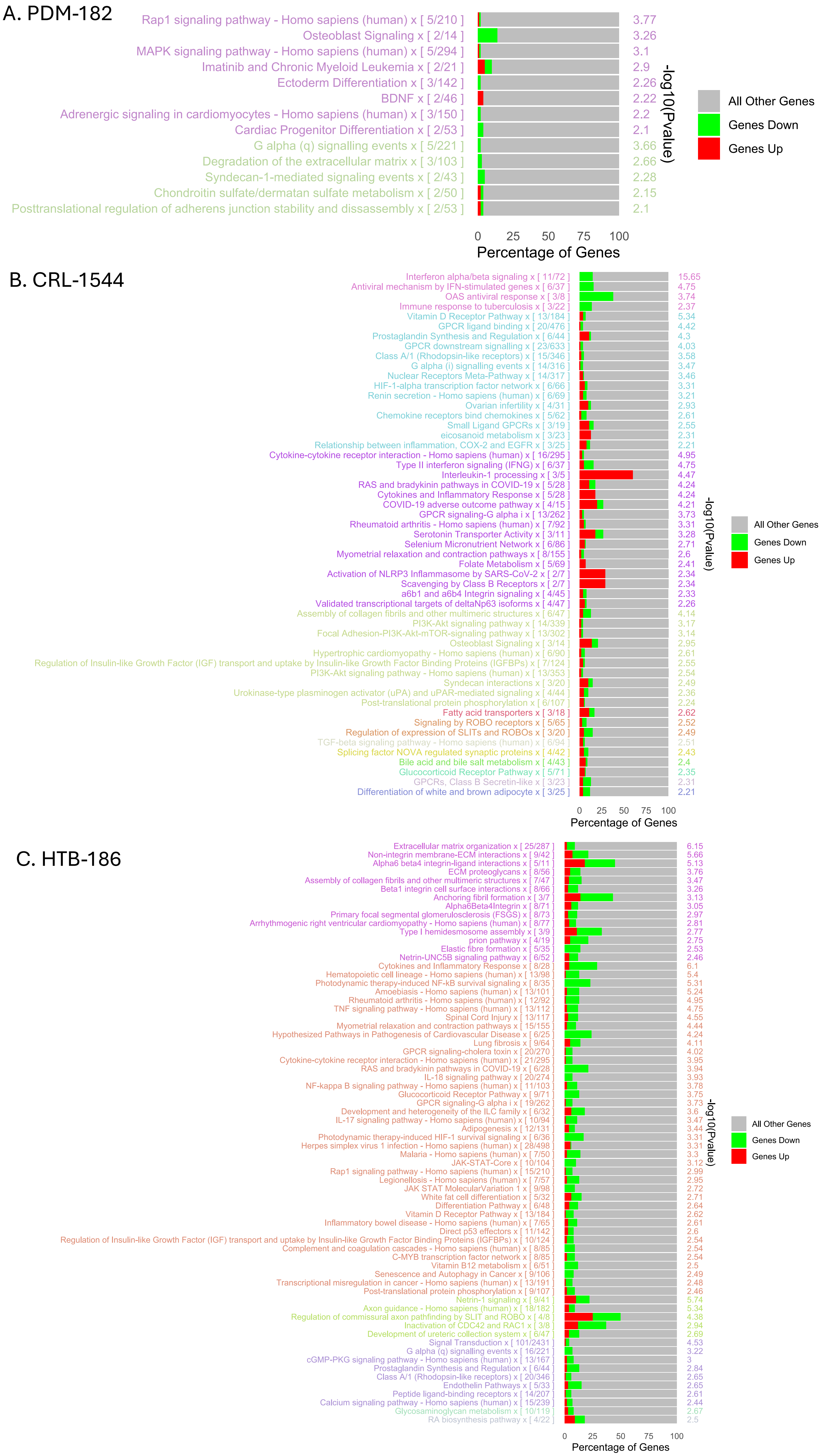

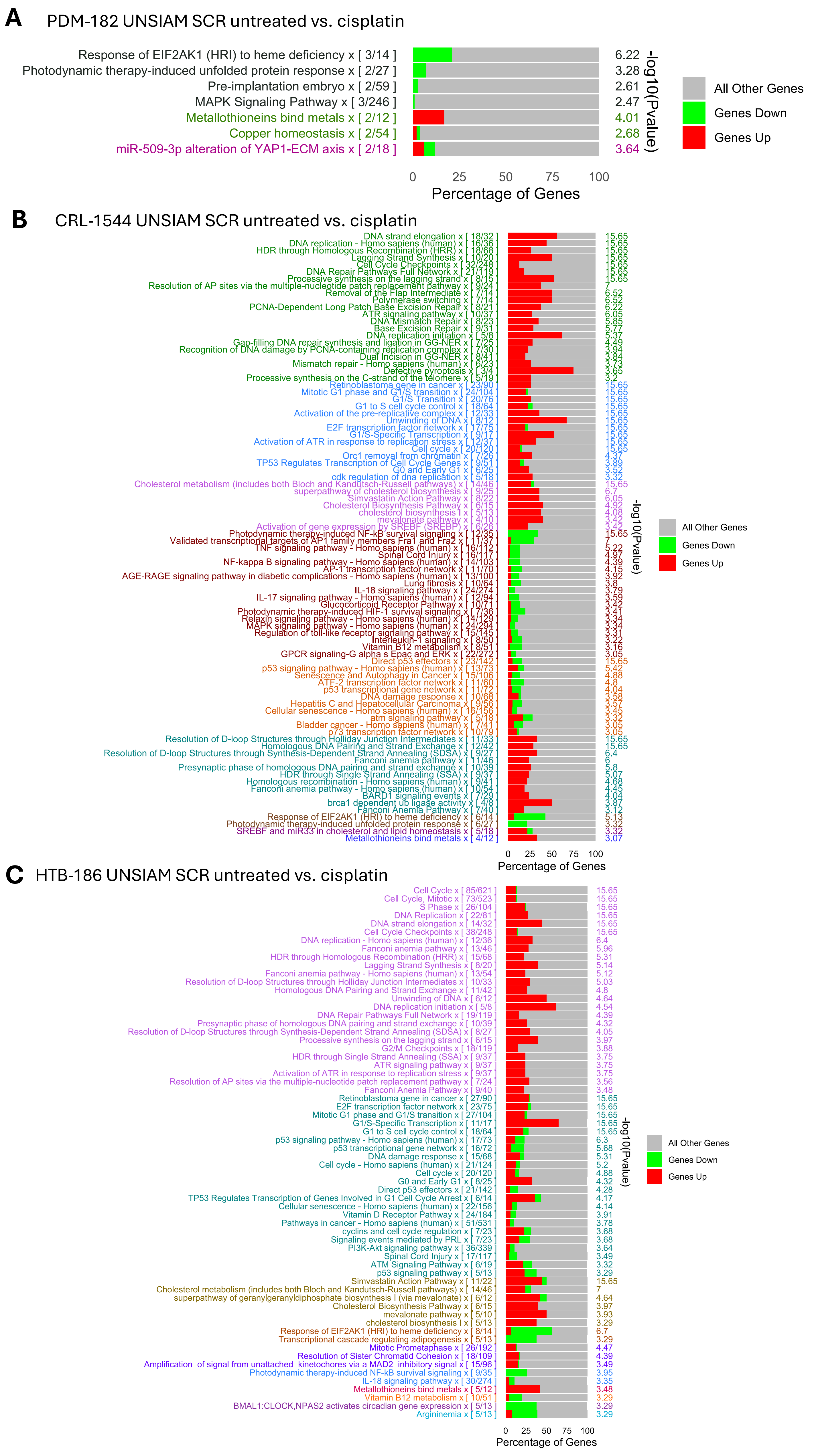

A
PDM-182 UNSIAM SCR untreated vs. cisplatin
Response of EIF2AK1 (HRI) to heme deficiency x [ 3/14 ] 6.22
Photodynamic therapy-induced unfolded protein response x [ 2/27 ] 3.28
Pre-implantation embryo x [ 2/59 ] 2.61
MAPK Signaling Pathway x [ 3/246 ] 2.47
Metallothioneins bind metals x [ 2/12 ] 4.01
Copper homeostasis x [ 2/54 ] 2.68
miR-509-3p alteration of YAP1-ECM axis x [ 2/18 ] 3.64
0 25 50 75 100
Percentage of Genes
-log10(Pvalue)
All Other Genes
Genes Down
Genes Up
B
CRL-1544 UNSIAM SCR untreated vs. cisplatin
DNA strand elongation x [ 18/32 ] 15.65
DNA replication - Homo sapiens (human) x [ 16/36 ] 15.65
HDR through Homologous Recombination (HRR) x [ 18/68 ] 15.65
Lagging Strand Synthesis x [ 10/20 ] 15.65
Cell Cycle Checkpoints x [ 32/248 ] 15.65
DNA Repair Pathways Full Network x [ 21/119 ] 15.65
Processive synthesis on the lagging strand x [ 8/15 ] 15.65
Resolution of AP sites via the multiple-nucleotide patch replacement pathway x [ 9/24 ] 7
Removal of the Flap Intermediate x [ 7/14 ] 6.52
Polymerase switching x [ 7/14 ] 6.52
PCNA-Dependent Long Patch Base Excision Repair x [ 8/21 ] 6.22
ATR signaling pathway x [ 10/37 ] 6.05
DNA Mismatch Repair x [ 8/23 ] 5.85
Base Excision Repair x [ 9/31 ] 5.77
DNA replication initiation x [ 5/8 ] 5.37
Gap-filling DNA repair synthesis and ligation in GG-NER x [ 7/25 ] 4.49
Recognition of DNA damage by PCNA-containing replication complex x [ 7/30 ] 3.94
Dual Incision in GG-NER x [ 8/41 ] 3.84
Mismatch repair - Homo sapiens (human) x [ 6/23 ] 3.73
Defective pyroptosis x [ 3/4 ] 3.65
Processive synthesis on the C-strand of the telomere x [ 5/19 ] 3.2
Retinoblastoma gene in cancer x [ 23/90 ] 15.65
Mitotic G1 phase and G1/S transition x [ 24/104 ] 15.65
G1/S Transition x [ 20/76 ] 15.65
G1 to S cell cycle control x [ 18/64 ] 15.65
Activation of the pre-replicative complex x [ 12/33 ] 15.65
Unwinding of DNA x [ 8/12 ] 15.65
E2F transcription factor network x [ 17/75 ] 15.65
G1/S-Specific Transcription x [ 9/17 ] 15.65
Activation of ATR in response to replication stress x [ 12/37 ] 15.65
Cell cycle x [ 20/120 ] 15.65
Orc1 removal from chromatin x [ 7/26 ] 4.37
TP53 Regulates Transcription of Cell Cycle Genes x [ 9/51 ] 3.89
G0 and Early G1 x [ 6/25 ] 3.52
cdk regulation of dna replication x [ 5/18 ] 3.32
Cholesterol metabolism (includes both Bloch and Kandutsch-Russell pathways) x [ 14/46 ] 15.65
superpathway of cholesterol biosynthesis x [ 9/25 ] 6.7
Simvastatin Action Pathway x [ 8/22 ] 6.05
Cholesterol Biosynthesis Pathway x [ 6/15 ] 4.92
cholesterol biosynthesis I x [ 5/13 ] 4.08
mevalonate pathway x [ 4/10 ] 3.42
Activation of gene expression by SREBF (SREBP) x [ 6/26 ] 3.42
Photodynamic therapy-induced NF-kB survival signaling x [ 12/35 ] 15.65
Validated transcriptional targets of AP1 family members Fra1 and Fra2 x [ 11/37 ] 7
TNF signaling pathway - Homo sapiens (human) x [ 16/112 ] 5.22
Spinal Cord Injury x [ 16/117 ] 4.97
NF-kappa B signaling pathway - Homo sapiens (human) x [ 14/103 ] 4.39
AP-1 transcription factor network x [ 11/70 ] 4.15
AGE-RAGE signaling pathway in diabetic complications - Homo sapiens (human) x [ 13/100 ] 3.92
Lung fibrosis x [ 10/64 ] 3.8
IL-18 signaling pathway x [ 24/274 ] 3.79
IL-17 signaling pathway - Homo sapiens (human) x [ 12/94 ] 3.59
Glucocorticoid Receptor Pathway x [ 10/71 ] 3.42
Photodynamic therapy-induced HIF-1 survival signaling x [ 7/36 ] 3.41
Relaxin signaling pathway - Homo sapiens (human) x [ 14/129 ] 3.34
MAPK signaling pathway - Homo sapiens (human) x [ 24/294 ] 3.34
Regulation of toll-like receptor signaling pathway x [ 15/145 ] 3.31
Interleukin-1 signaling x [ 8/50 ] 3.22
Vitamin B12 metabolism x [ 8/51 ] 3.16
GPCR signaling-G alpha s Epac and ERK x [ 22/272 ] 3.05
Direct p53 effectors x [ 23/142 ] 15.65
p53 signaling pathway - Homo sapiens (human) x [ 13/73 ] 5.42
Senescence and Autophagy in Cancer x [ 15/106 ] 4.88
ATF-2 transcription factor network x [ 11/60 ] 4.8
p53 transcriptional gene network x [ 11/72 ] 4.04
DNA damage response x [ 10/68 ] 3.58
Hepatitis C and Hepatocellular Carcinoma x [ 9/56 ] 3.57
Cellular senescence - Homo sapiens (human) x [ 16/156 ] 3.45
atm signaling pathway x [ 5/18 ] 3.32
Bladder cancer - Homo sapiens (human) x [ 7/41 ] 3.05
p73 transcription factor network x [ 10/79 ] 3.05
Resolution of D-loop Structures through Holliday Junction Intermediates x [ 11/33 ] 15.65
Homologous DNA Pairing and Strand Exchange x [ 12/42 ] 15.65
Resolution of D-loop Structures through Synthesis-Dependent Strand Annealing (SDSA) x [ 9/27 ] 6.4
Fanconi anemia pathway x [ 11/46 ] 6
Presynaptic phase of homologous DNA pairing and strand exchange x [ 10/39 ] 5.8
HDR through Single Strand Annealing (SSA) x [ 9/37 ] 5.07
Homologous recombination - Homo sapiens (human) x [ 9/41 ] 4.68
Fanconi anemia pathway - Homo sapiens (human) x [ 10/54 ] 4.45
BARD1 signaling events x [ 7/29 ] 4.04
brca1 dependent ub ligase activity x [ 4/8 ] 3.87
Fanconi Anemia Pathway x [ 7/40 ] 3.12
Response of EIF2AK1 (HRI) to heme deficiency x [ 6/14 ] 5.13
Photodynamic therapy-induced unfolded protein response x [ 6/27 ] 3.32
SREBF and miR33 in cholesterol and lipid homeostasis x [ 5/18 ] 3.32
Metallothioneins bind metals x [ 4/12 ] 3.07
0 25 50 75 100
Percentage of Genes
-log10(Pvalue)
All Other Genes
Genes Down
Genes Up
C
HTB-186 UNSIAM SCR untreated vs. cisplatin
Cell Cycle x [ 85/621 ] 15.65
Cell Cycle, Mitotic x [ 73/523 ] 15.65
S Phase x [ 26/104 ] 15.65
DNA Replication x [ 22/81 ] 15.65
DNA strand elongation x [ 14/32 ] 15.65
Cell Cycle Checkpoints x [ 38/248 ] 15.65
DNA replication - Homo sapiens (human) x [ 12/36 ] 6.4
Fanconi anemia pathway x [ 13/46 ] 5.96
HDR through Homologous Recombination (HRR) x [ 15/68 ] 5.31
Lagging Strand Synthesis x [ 8/20 ] 5.14
Fanconi anemia pathway - Homo sapiens (human) x [ 13/54 ] 5.12
Resolution of D-loop Structures through Holliday Junction Intermediates x [ 10/33 ] 5.03
Homologous DNA Pairing and Strand Exchange x [ 11/42 ] 4.8
Unwinding of DNA x [ 6/12 ] 4.64
DNA replication initiation x [ 5/8 ] 4.54
DNA Repair Pathways Full Network x [ 19/119 ] 4.39
Presynaptic phase of homologous DNA pairing and strand exchange x [ 10/39 ] 4.32
Resolution of D-loop Structures through Synthesis-Dependent Strand Annealing (SDSA) x [ 8/27 ] 4.05
Processive synthesis on the lagging strand x [ 6/15 ] 3.97
G2/M Checkpoints x [ 18/119 ] 3.88
HDR through Single Strand Annealing (SSA) x [ 9/37 ] 3.75
ATR signaling pathway x [ 9/37 ] 3.75
Activation of ATR in response to replication stress x [ 9/37 ] 3.75
Resolution of AP sites via the multiple-nucleotide patch replacement pathway x [ 7/24 ] 3.56
Fanconi Anemia Pathway x [ 9/40 ] 3.48
Retinoblastoma gene in cancer x [ 27/90 ] 15.65
E2F transcription factor network x [ 23/75 ] 15.65
Mitotic G1 phase and G1/S transition x [ 27/104 ] 15.65
G1/S-Specific Transcription x [ 11/17 ] 15.65
G1 to S cell cycle control x [ 18/64 ] 15.65
p53 signaling pathway - Homo sapiens (human) x [ 17/73 ] 6.3
p53 transcriptional gene network x [ 16/72 ] 5.68
DNA damage response x [ 15/68 ] 5.31
Cell cycle - Homo sapiens (human) x [ 21/124 ] 5.2
Cell cycle x [ 20/120 ] 4.88
G0 and Early G1 x [ 8/25 ] 4.32
Direct p53 effectors x [ 21/142 ] 4.28
TP53 Regulates Transcription of Genes Involved in G1 Cell Cycle Arrest x [ 6/14 ] 4.17
Cellular senescence - Homo sapiens (human) x [ 22/156 ] 4.14
Vitamin D Receptor Pathway x [ 24/184 ] 3.91
Pathways in cancer - Homo sapiens (human) x [ 51/531 ] 3.78
cyclins and cell cycle regulation x [ 7/23 ] 3.68
Signaling events mediated by PRL x [ 7/23 ] 3.68
PI3K-Akt signaling pathway x [ 36/339 ] 3.64
Spinal Cord Injury x [ 17/117 ] 3.49
ATM Signaling Pathway x [ 6/19 ] 3.32
p53 signaling pathway x [ 5/13 ] 3.29
Simvastatin Action Pathway x [ 11/22 ] 15.65
Cholesterol metabolism (includes both Bloch and Kandutsch-Russell pathways) x [ 14/46 ] 7
superpathway of geranylgeranyldiphosphate biosynthesis I (via mevalonate) x [ 6/12 ] 4.64
Cholesterol Biosynthesis Pathway x [ 6/15 ] 3.97
mevalonate pathway x [ 5/10 ] 3.93
cholesterol biosynthesis I x [ 5/13 ] 3.29
Response of EIF2AK1 (HRI) to heme deficiency x [ 8/14 ] 6.7
Transcriptional cascade regulating adipogenesis x [ 5/13 ] 3.29
Mitotic Prometaphase x [ 26/192 ] 4.47
Resolution of Sister Chromatid Cohesion x [ 18/109 ] 4.39
Amplification of signal from unattached kinetochores via a MAD2 inhibitory signal x [ 15/96 ] 3.49
Photodynamic therapy-induced NF-kB survival signaling x [ 9/35 ] 3.95
IL-18 signaling pathway x [ 30/274 ] 3.35
Metallothioneins bind metals x [ 5/12 ] 3.48
Vitamin B12 metabolism x [ 10/51 ] 3.29
BMAL1:CLOCK,NPAS2 activates circadian gene expression x [ 5/13 ] 3.29
Argininemia x [ 5/13 ] 3.29
0 25 50 75 100
Percentage of Genes
-log10(Pvalue)
All Other Genes
Genes Down
Genes Up

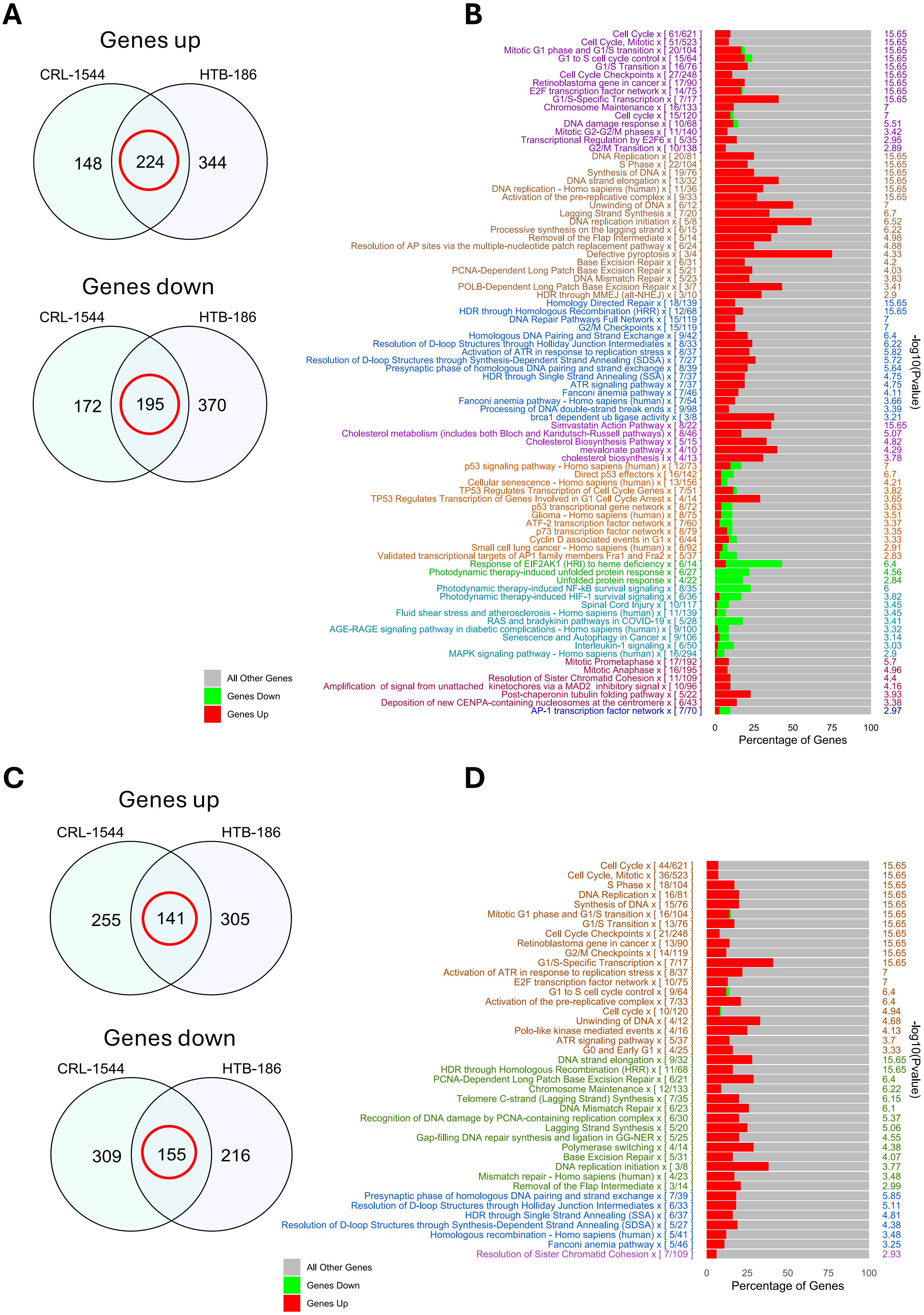
A
Genes up
CRL-1544
HTB-186
148
224
344
Genes down
CRL-1544
HTB-186
172
195
370
All Other Genes
Genes Down
Genes Up
B
Cell Cycle x [ 61/621 ] 15.65
Cell Cycle, Mitotic x [ 51/523 ] 15.65
Mitotic G1 phase and G1/S transition x [ 20/104 ] 15.65
G1 to S cell cycle control x [ 15/64 ] 15.65
G1/S Transition x [ 16/76 ] 15.65
Cell Cycle Checkpoints x [ 27/248 ] 15.65
Retinoblastoma gene in cancer x [ 17/90 ] 15.65
E2F transcription factor network x [ 14/75 ] 15.65
G1/S-Specific Transcription x [ 7/17 ] 15.65
Chromosome Maintenance x [ 16/133 ] 7
Cell cycle x [ 15/120 ] 7
DNA damage response x [ 10/68 ] 5.51
Mitotic G2-G2/M phases x [ 11/140 ] 3.42
Transcriptional Regulation by E2F6 x [ 5/35 ] 2.95
G2/M Transition x [ 10/138 ] 2.89
DNA Replication x [ 20/81 ] 15.65
S Phase x [ 22/104 ] 15.65
Synthesis of DNA x [ 19/76 ] 15.65
DNA strand elongation x [ 13/32 ] 15.65
DNA replication - Homo sapiens (human) x [ 11/36 ] 15.65
Activation of the pre-replicative complex x [ 9/33 ] 15.65
Unwinding of DNA x [ 6/12 ] 7
Lagging Strand Synthesis x [ 7/20 ] 6.7
DNA replication initiation x [ 5/8 ] 6.52
Processive synthesis on the lagging strand x [ 6/15 ] 6.22
Removal of the Flap Intermediate x [ 5/14 ] 4.98
Resolution of AP sites via the multiple-nucleotide patch replacement pathway x [ 6/24 ] 4.88
Defective pyroptosis x [ 3/4 ] 4.33
Base Excision Repair x [ 6/31 ] 4.2
PCNA-Dependent Long Patch Base Excision Repair x [ 5/21 ] 4.03
DNA Mismatch Repair x [ 5/23 ] 3.83
POLB-Dependent Long Patch Base Excision Repair x [ 3/7 ] 3.41
HDR through MMEJ (alt-NHEJ) x [ 3/10 ] 2.9
Homology Directed Repair x [ 18/139 ] 15.65
HDR through Homologous Recombination (HRR) x [ 12/68 ] 15.65
DNA Repair Pathways Full Network x [ 15/119 ] 7
G2/M Checkpoints x [ 15/119 ] 7
Homologous DNA Pairing and Strand Exchange x [ 9/42 ] 6.4
Resolution of D-loop Structures through Holliday Junction Intermediates x [ 8/33 ] 6.22
Activation of ATR in response to replication stress x [ 8/37 ] 5.82
Resolution of D-loop Structures through Synthesis-Dependent Strand Annealing (SDSA) x [ 7/27 ] 5.72
Presynaptic phase of homologous DNA pairing and strand exchange x [ 8/39 ] 5.64
HDR through Single Strand Annealing (SSA) x [ 7/37 ] 4.75
ATR signaling pathway x [ 7/37 ] 4.75
Fanconi anemia pathway x [ 7/46 ] 4.11
Fanconi anemia pathway - Homo sapiens (human) x [ 7/54 ] 3.66
Processing of DNA double-strand break ends x [ 9/98 ] 3.39
brca1 dependent ub ligase activity x [ 3/8 ] 3.21
Simvastatin Action Pathway x [ 8/22 ] 15.65
Cholesterol metabolism (includes both Bloch and Kandutsch-Russell pathways) x [ 8/46 ] 5.07
Cholesterol Biosynthesis Pathway x [ 5/15 ] 4.82
mevalonate pathway x [ 4/10 ] 4.29
cholesterol biosynthesis I x [ 4/13 ] 3.78
p53 signaling pathway - Homo sapiens (human) x [ 12/73 ] 7
Direct p53 effectors x [ 16/142 ] 6.7
Cellular senescence - Homo sapiens (human) x [ 13/156 ] 4.21
TP53 Regulates Transcription of Cell Cycle Genes x [ 7/51 ] 3.82
TP53 Regulates Transcription of Genes Involved in G1 Cell Cycle Arrest x [ 4/14 ] 3.65
p53 transcriptional gene network x [ 8/72 ] 3.63
Glioma - Homo sapiens (human) x [ 8/75 ] 3.51
ATF-2 transcription factor network x [ 7/60 ] 3.37
p73 transcription factor network x [ 8/79 ] 3.35
Cyclin D associated events in G1 x [ 6/44 ] 3.33
Small cell lung cancer - Homo sapiens (human) x [ 8/92 ] 2.91
Validated transcriptional targets of AP1 family members Fra1 and Fra2 x [ 5/37 ] 2.83
Response of EIF2AK1 (HRI) to heme deficiency x [ 6/14 ] 6.4
Photodynamic therapy-induced unfolded protein response x [ 6/27 ] 4.56
Unfolded protein response x [ 4/22 ] 2.84
Photodynamic therapy-induced NF-kB survival signaling x [ 8/35 ] 6
Photodynamic therapy-induced HIF-1 survival signaling x [ 6/36 ] 3.82
Spinal Cord Injury x [ 10/117 ] 3.45
Fluid shear stress and atherosclerosis - Homo sapiens (human) x [ 11/139 ] 3.45
RAS and bradykinin pathways in COVID-19 x [ 5/28 ] 3.41
AGE-RAGE signaling pathway in diabetic complications - Homo sapiens (human) x [ 9/100 ] 3.32
Senescence and Autophagy in Cancer x [ 9/106 ] 3.14
Interleukin-1 signaling x [ 6/50 ] 3.03
MAPK signaling pathway - Homo sapiens (human) x [ 16/294 ] 2.9
Mitotic Prometaphase x [ 17/192 ] 5.7
Mitotic Anaphase x [ 16/195 ] 4.96
Resolution of Sister Chromatid Cohesion x [ 11/109 ] 4.4
Amplification of signal from unattached kinetochores via a MAD2 inhibitory signal x [ 10/96 ] 4.16
Post-chaperonin tubulin folding pathway x [ 5/22 ] 3.93
Deposition of new CENPA-containing nucleosomes at the centromere x [ 6/43 ] 3.38
AP-1 transcription factor network x [ 7/70 ] 2.97
0 25 50 75 100
Percentage of Genes
-log10(Pvalue)
C
Genes up
CRL-1544
HTB-186
255
141
305
Genes down
CRL-1544
HTB-186
309
155
216
All Other Genes
Genes Down
Genes Up
D
Cell Cycle x [ 44/621 ] 15.65
Cell Cycle, Mitotic x [ 36/523 ] 15.65
S Phase x [ 18/104 ] 15.65
DNA Replication x [ 16/81 ] 15.65
Synthesis of DNA x [ 15/76 ] 15.65
Mitotic G1 phase and G1/S transition x [ 16/104 ] 15.65
G1/S Transition x [ 13/76 ] 15.65
Cell Cycle Checkpoints x [ 21/248 ] 15.65
Retinoblastoma gene in cancer x [ 13/90 ] 15.65
G2/M Checkpoints x [ 14/119 ] 15.65
G1/S-Specific Transcription x [ 7/17 ] 15.65
Activation of ATR in response to replication stress x [ 8/37 ] 7
E2F transcription factor network x [ 10/75 ] 7
G1 to S cell cycle control x [ 9/64 ] 6.4
Activation of the pre-replicative complex x [ 7/33 ] 6.4
Cell cycle x [ 10/120 ] 4.94
Unwinding of DNA x [ 4/12 ] 4.68
Polo-like kinase mediated events x [ 4/16 ] 4.13
ATR signaling pathway x [ 5/37 ] 3.7
G0 and Early G1 x [ 4/25 ] 3.33
DNA strand elongation x [ 9/32 ] 15.65
HDR through Homologous Recombination (HRR) x [ 11/68 ] 15.65
PCNA-Dependent Long Patch Base Excision Repair x [ 6/21 ] 6.4
Chromosome Maintenance x [ 12/133 ] 6.22
Telomere C-strand (Lagging Strand) Synthesis x [ 7/35 ] 6.15
DNA Mismatch Repair x [ 6/23 ] 6.1
Recognition of DNA damage by PCNA-containing replication complex x [ 6/30 ] 5.37
Lagging Strand Synthesis x [ 5/20 ] 5.06
Gap-filling DNA repair synthesis and ligation in GG-NER x [ 5/25 ] 4.55
Polymerase switching x [ 4/14 ] 4.38
Base Excision Repair x [ 5/31 ] 4.07
DNA replication initiation x [ 3/8 ] 3.77
Mismatch repair - Homo sapiens (human) x [ 4/23 ] 3.48
Removal of the Flap Intermediate x [ 3/14 ] 2.99
Presynaptic phase of homologous DNA pairing and strand exchange x [ 7/39 ] 5.85
Resolution of D-loop Structures through Holliday Junction Intermediates x [ 6/33 ] 5.11
HDR through Single Strand Annealing (SSA) x [ 6/37 ] 4.81
Resolution of D-loop Structures through Synthesis-Dependent Strand Annealing (SDSA) x [ 5/27 ] 4.38
Homologous recombination - Homo sapiens (human) x [ 5/41 ] 3.48
Fanconi anemia pathway x [ 5/46 ] 3.25
Resolution of Sister Chromatid Cohesion x [ 7/109 ] 2.93
0 25 50 75 100
Percentage of Genes
-log10(Pvalue)

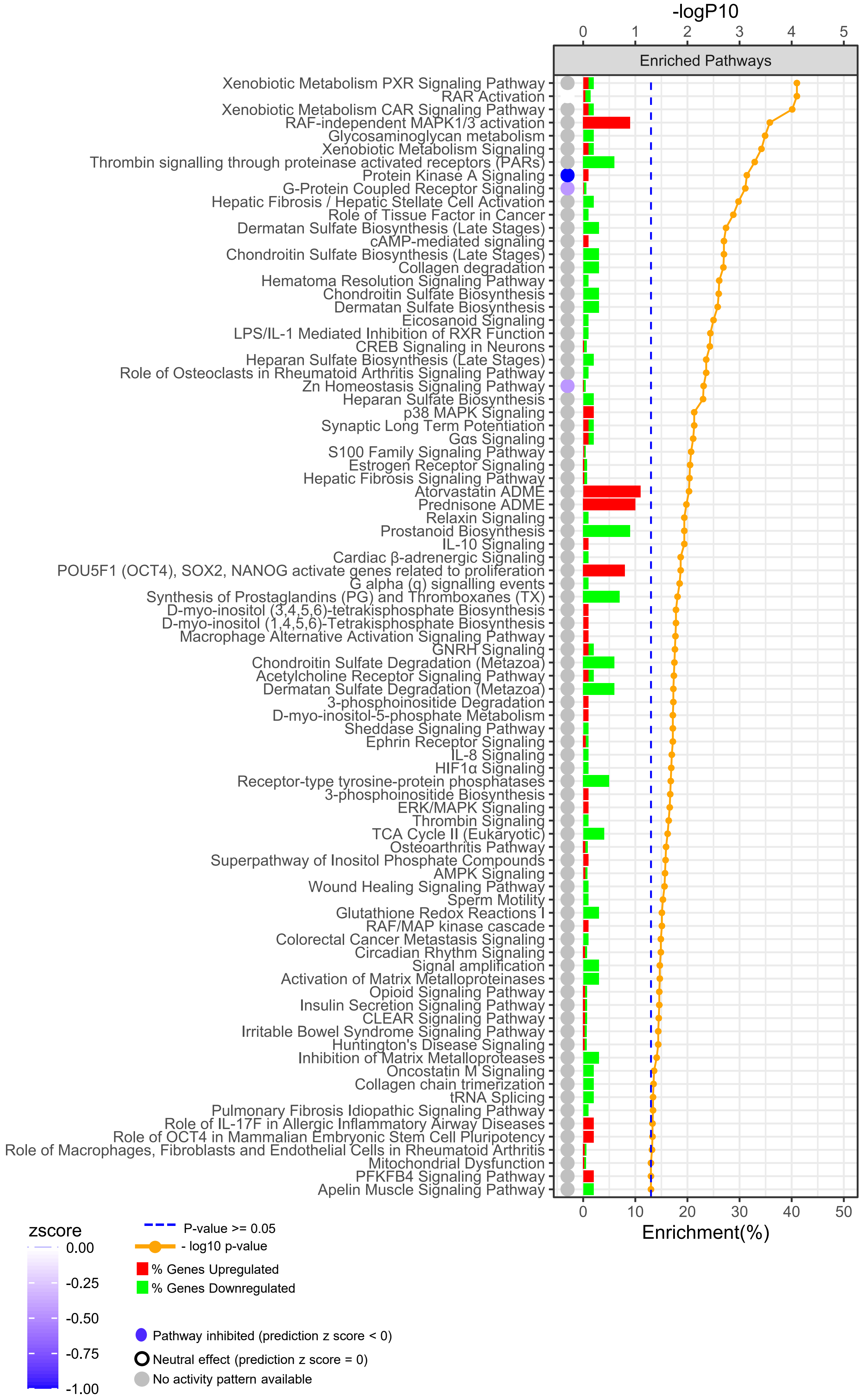

-logP10
0
1
2
3
4
5
Enriched Pathways
Xenobiotic Metabolism PXR Signaling Pathway
RAR Activation
Xenobiotic Metabolism CAR Signaling Pathway
RAF-independent MAPK1/3 activation
Glycosaminoglycan metabolism
Xenobiotic Metabolism Signaling
Thrombin signalling through proteinase activated receptors (PARs)
Protein Kinase A Signaling
G-Protein Coupled Receptor Signaling
Hepatic Fibrosis / Hepatic Stellate Cell Activation
Role of Tissue Factor in Cancer
Dermatan Sulfate Biosynthesis (Late Stages)
cAMP-mediated signaling
Chondroitin Sulfate Biosynthesis (Late Stages)
Collagen degradation
Hematoma Resolution Signaling Pathway
Chondroitin Sulfate Biosynthesis
Dermatan Sulfate Biosynthesis
Eicosanoid Signaling
LPS/IL-1 Mediated Inhibition of RXR Function
CREB Signaling in Neurons
Heparan Sulfate Biosynthesis (Late Stages)
Role of Osteoclasts in Rheumatoid Arthritis Signaling Pathway
Zn Homeostasis Signaling Pathway
Heparan Sulfate Biosynthesis
p38 MAPK Signaling
Synaptic Long Term Potentiation
Gαs Signaling
S100 Family Signaling Pathway
Estrogen Receptor Signaling
Hepatic Fibrosis Signaling Pathway
Atorvastatin ADME
Prednisone ADME
Relaxin Signaling
Prostanoid Biosynthesis
IL-10 Signaling
Cardiac β-adrenergic Signaling
POU5F1 (OCT4), SOX2, NANOG activate genes related to proliferation
G alpha (q) signalling events
Synthesis of Prostaglandins (PG) and Thromboxanes (TX)
D-myo-inositol (3,4,5,6)-tetrakisphosphate Biosynthesis
D-myo-inositol (1,4,5,6)-Tetrakisphosphate Biosynthesis
Macrophage Alternative Activation Signaling Pathway
GNRH Signaling
Chondroitin Sulfate Degradation (Metazoa)
Acetylcholine Receptor Signaling Pathway
Dermatan Sulfate Degradation (Metazoa)
3-phosphoinositide Degradation
D-myo-inositol-5-phosphate Metabolism
Sheddase Signaling Pathway
Ephrin Receptor Signaling
IL-8 Signaling
HIF1α Signaling
Receptor-type tyrosine-protein phosphatases
3-phosphoinositide Biosynthesis
ERK/MAPK Signaling
Thrombin Signaling
TCA Cycle II (Eukaryotic)
Osteoarthritis Pathway
Superpathway of Inositol Phosphate Compounds
AMPK Signaling
Wound Healing Signaling Pathway
Sperm Motility
Glutathione Redox Reactions I
RAF/MAP kinase cascade
Colorectal Cancer Metastasis Signaling
Circadian Rhythm Signaling
Signal amplification
Activation of Matrix Metalloproteinases
Opioid Signaling Pathway
Insulin Secretion Signaling Pathway
CLEAR Signaling Pathway
Irritable Bowel Syndrome Signaling Pathway
Huntington's Disease Signaling
Inhibition of Matrix Metalloproteases
Oncostatin M Signaling
Collagen chain trimerization
tRNA Splicing
Pulmonary Fibrosis Idiopathic Signaling Pathway
Role of IL-17F in Allergic Inflammatory Airway Diseases
Role of OCT4 in Mammalian Embryonic Stem Cell Pluripotency
Role of Macrophages, Fibroblasts and Endothelial Cells in Rheumatoid Arthritis
Mitochondrial Dysfunction
PFKFB4 Signaling Pathway
Apelin Muscle Signaling Pathway
0
10
20
30
40
50
Enrichment(%)
zscore
0.00
-0.25
-0.50
-0.75
-1.00
P-value >= 0.05
- log10 p-value
% Genes Upregulated
% Genes Downregulated
Pathway inhibited (prediction z score < 0)
Neutral effect (prediction z score = 0)
No activity pattern available

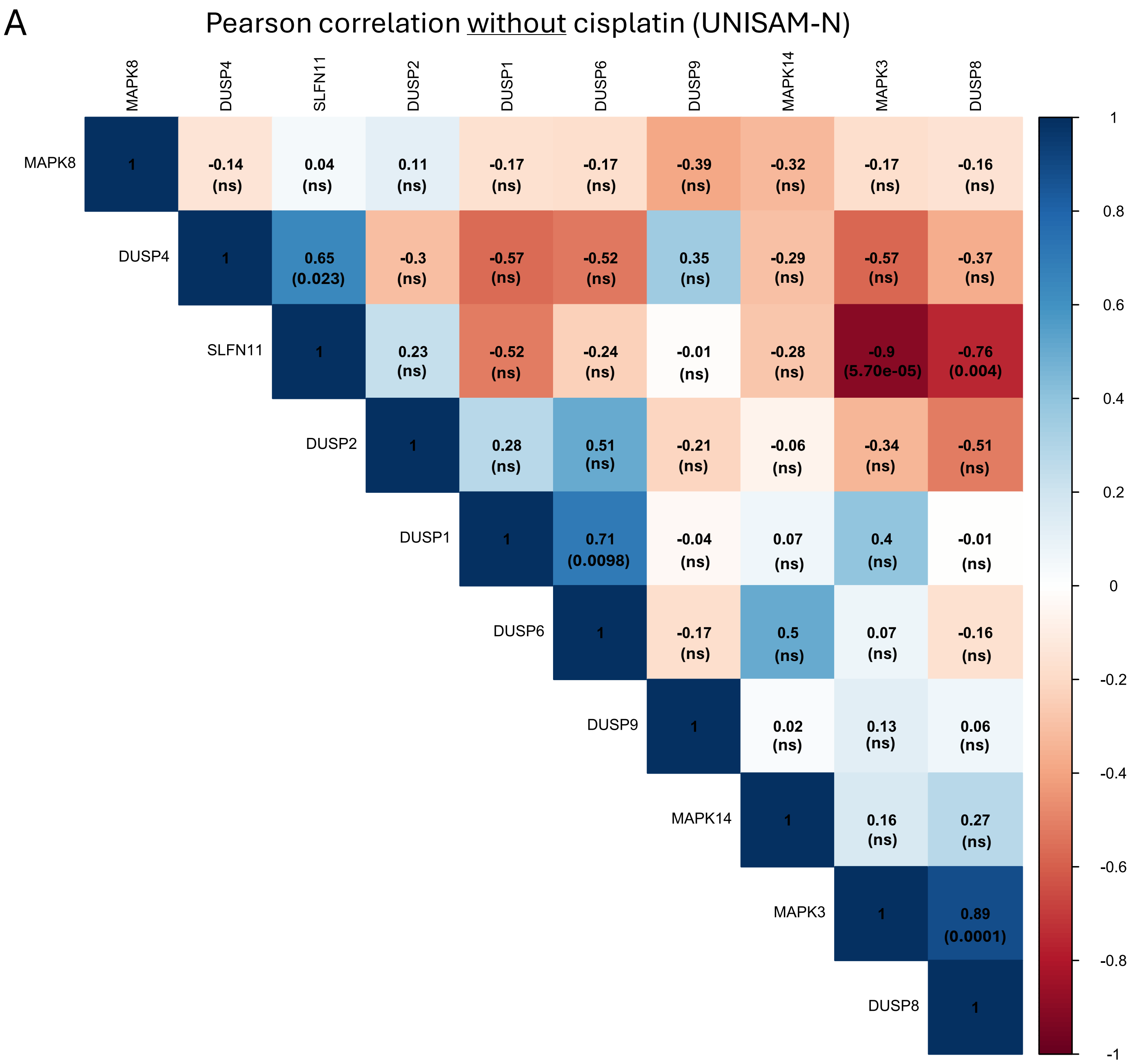
A
Pearson correlation without cisplatin (UNISAM-N)
MAPK8
DUSP4
SLFN11
DUSP2
DUSP1
DUSP6
DUSP9
MAPK14
MAPK3
DUSP8
1 -0.14 (ns) 0.04 (ns) 0.11 (ns) -0.17 (ns) -0.17 (ns) -0.39 (ns) -0.32 (ns) -0.17 (ns) -0.16 (ns)
1 0.65 (0.023) -0.3 (ns) -0.57 (ns) -0.52 (ns) 0.35 (ns) -0.29 (ns) -0.57 (ns) -0.37 (ns)
1 0.23 (ns) -0.52 (ns) -0.24 (ns) -0.01 (ns) -0.28 (ns) -0.9 (5.70e-05) -0.76 (0.004)
1 0.28 (ns) 0.51 (ns) -0.21 (ns) -0.06 (ns) -0.34 (ns) -0.51 (ns)
1 0.71 (0.0098) -0.04 (ns) 0.07 (ns) 0.4 (ns) -0.01 (ns)
1 -0.17 (ns) 0.5 (ns) 0.07 (ns) -0.16 (ns)
1 0.02 (ns) 0.13 (ns) 0.06 (ns)
1 0.16 (ns) 0.27 (ns)
1 0.89 (0.0001)
1
1
0.8
0.6
0.4
0.2
0
-0.2
-0.4
-0.6
-0.8
-1

A Regulation of map kinase pathway by DUSPs

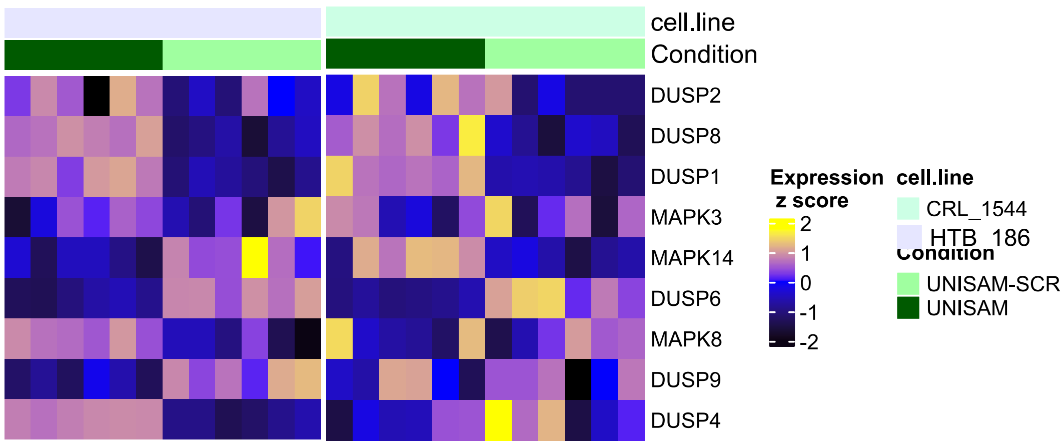


**B** Pearson correlation with cisplatin (UNISAM-N)

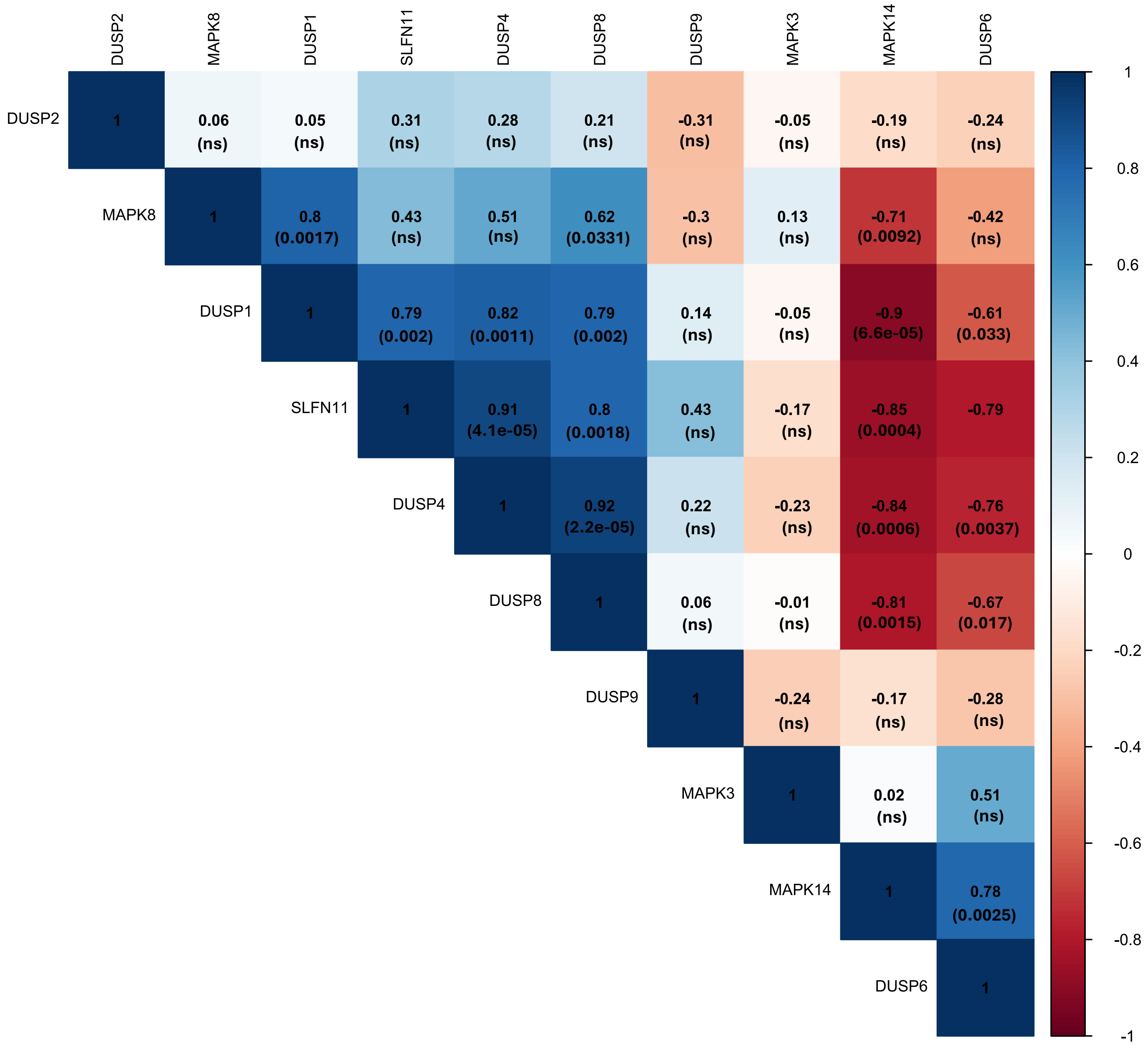
